# Haplotype-Resolved Cattle Genomes Provide Insights Into Structural Variation and Adaptation

**DOI:** 10.1101/720797

**Authors:** Wai Yee Low, Rick Tearle, Ruijie Liu, Sergey Koren, Arang Rhie, Derek M. Bickhart, Benjamin D. Rosen, Zev N. Kronenberg, Sarah B. Kingan, Elizabeth Tseng, Françoise Thibaud-Nissen, Fergal J. Martin, Konstantinos Billis, Jay Ghurye, Alex R. Hastie, Joyce Lee, Andy W.C. Pang, Michael P. Heaton, Adam M. Phillippy, Stefan Hiendleder, Timothy P.L. Smith, John L. Williams

**Affiliations:** The Davies Research Centre, School of Animal and Veterinary Sciences, University of Adelaide, Roseworthy, SA 5371, Australia; Genome Informatics Section, Computational and Statistical Genomics Branch, National Human Genome Research Institute, Bethesda, Maryland, USA; Dairy Forage Research Center, ARS USDA, Madison, Wisconsin, USA; Animal Genomics and Improvement Laboratory, ARS USDA, Beltsville, Maryland, USA; Phase Genomics, 4000 Mason Road, Suite 225, Seattle, WA 98195, USA; Pacific Biosciences, Menlo Park, CA 94025, USA; National Center for Biotechnology Information, National Library of Medicine, National Institutes of Health, Bethesda, MD 20894, USA; European Molecular Biology Laboratory, European Bioinformatics Institute, Wellcome Genome Campus, Hinxton, Cambridge CB10 1SD, UK; Center for Bioinformatics and Computational Biology, Lab 3104A, Biomolecular Science Building, University of Maryland, College Park, Maryland - 20742; BioNano Genomics, San Diego, California, USA; US Meat Animal Research Center, ARS USDA, Clay Center, Nebraska, USA

## Abstract

We present high quality, phased genome assemblies representative of taurine and indicine cattle, subspecies that differ markedly in productivity-related traits and environmental adaptation. We report a new haplotype-aware scaffolding and polishing pipeline using contigs generated by the trio binning method to produce haplotype-resolved, chromosome-level genome assemblies of Angus (taurine) and Brahman (indicine) cattle breeds. These assemblies were used to identify structural and copy number variants that differentiate the subspecies and we found variant detection was sensitive to the specific reference genome chosen. Six gene families with immune related functions are expanded in the indicine lineage. Assembly of the genomes of both subspecies from a single individual enabled transcripts to be phased to detect allele-specific expression, and to study genome-wide selective sweeps. An indicus-specific extra copy of fatty acid desaturase is under positive selection and may contribute to indicine adaptation to heat and drought.

## Main

About 10,000 years ago, cattle were domesticated from the aurochs which ranged across Eurasia and North Africa but are now extinct^1^. Modern day cattle belong to two subspecies, the humped zebu or indicine breeds (*Bos taurus indicus*) and the humpless taurine breeds (*Bos taurus taurus*), which arose from independent domestication events of genetically distinct aurochs populations^2^.

During the last century, taurine breeds have been intensively selected for production traits, particularly milk and meat yield, and generally have higher fertility than indicine breeds. European taurine breeds, such as Angus, have excellent carcass and meat quality, high fertility, and reach puberty early. These breeds have been imported by farmers around the world to improve or replace less-productive breeds. However, while European taurine animals are well adapted to temperate environments, they do not thrive in hot, humid tropical environments with high disease and parasite challenge.

Indicine breeds originated from the Indus valley and later spread to Africa and across southeast Asia^3^. Between 1854 and 1926, the four indicine breeds, Ongole, Krishna, Gir and Gujarat, were imported into the United States and crossed with European taurine cattle to create the Brahman breed. Current US Brahman cattle retain ∼10% of their genome of taurine origin^4^. Brahman have a short, thick, glossy coat that reflects sunlight and loose skin that increases the body surface area exposed for cooling. While the Brahman are less productive and have lower fertility than taurine breeds, they have desirable traits, such as heat tolerance, lower susceptibility to parasites such as ticks, and are more disease and drought resistant^5^.

We previously demonstrated a novel trio binning approach to assemble haplotypes of diploid individuals at the contig level. The quality of the contigs exceeded those of the best livestock reference genomes^6^. Here we present chromosome-level taurine (Angus) and indicine (Brahman) cattle genomes from a single crossbred individual that were assembled with haplotype-aware methodology that is less laborious than sequencing haploid clones^7^. The contiguity and accuracy of the final haplotype-resolved cattle assemblies set a new standard for diploid genomes and enable precise identification of genetic variants, from single nucleotide polymorphisms (SNPs) to large structural variants (SVs). A further benefit of haplotype-resolved genomes is that they can be used to better interpret allele-specific expression in diploid transcriptome profiles. Here we identify allele-specific and novel transcripts using PacBio Iso-Seq reads mapped onto the haplotype-resolved genomes. Considering the large differences in production and adaptation traits between taurine and indicine cattle, the genomes presented here are a milestone on the roadmap to identifying the molecular basis of globally important phenotypic traits that will help secure cattle production in a rapidly changing environment.

## Results

### *De novo* assembly and annotation of Angus and Brahman cattle genomes

The initial creation of haplotigs (haplotype-specific contigs) was presented in the description of the trio binning method implemented in TrioCanu^6^. Briefly, a male *Bos taurus* hybrid fetus, from an Angus sire and a Brahman dam, was sequenced to ∼136x long-read coverage, and the reads were sorted into parental haplotype bins based on unique sequence identified by short read sequencing of the parents prior to assembly with TrioCanu. The initial assemblies comprised 1747 Angus haplotigs and 1585 Brahman haplotigs (Table 1). The haplotig N50 was 29.4 Mb and 23.4 Mb for the Angus and Brahman, respectively.

**Table 1:** Assembly statistics

| Assembly | Software | Assembly level | Number of sequences<sup>a</sup> | Number of gaps | N50 (Mb) | Assembly size (Gb) |
| --- | --- | --- | --- | --- | --- | --- |
| **Angus** |  |  |  |  |  |  |
| PacBio | CANU | haplotig | 1747 | 0 | 29.4 | 2.6 |
| PacBio + Hi-C | SALSA2 | scaffold | 1515 | 235 | 104.6 | 2.6 |
| PacBio + Optical map | Bionano Access | scaffold | 1595 | 181 | 35.2 | 2.6 |
| UOA_Angus_1 | PBJelly, Aarow, custom scripts | chromosome | 1435 | 277 | 102.8 | 2.6 |
| **Brahman** |  |  |  |  |  |  |
| PacBio | CANU | haplotig | 1585 | 0 | 23.4 | 2.7 |
| PacBio + Hi-C | SALSA2 | scaffold | 1370 | 216 | 72.6 | 2.7 |
| PacBio + Optical map | Bionano Access | scaffold | 1353 | 268 | 31.7 | 2.7 |
| UOA_Brahman_1 | PBJelly, Aarow, custom scripts | chromosome | 1251 | 302 | 104.5 | 2.7 |

<sup>a</sup>There are 1,405 and 1,220 unplaced haplotigs in the final chromosome-level Angus and Brahman assemblies, respectively. These unplaced haplotigs comprise ~3.8% of total bases in the Angus assembly and ~2.1% of total bases in the Brahman assembly. Only the Brahman assembly has a complete mitochondrion sequence.

For the present study, additional data were generated for the same hybrid fetus, including ∼12x Hi-C reads, ∼167x Bionano optical map, and ∼84x Illumina paired-end reads (Fig. 1), to provide haplotype-resolved scaffolding and identify assembly errors. Following haplotig assembly, two sets of scaffolds, one based on Hi-C and the other on optical map data, were generated for each haplotype. Three different scaffolding programs (3D-DNA, Proximo and SALSA2) using the Hi-C data were evaluated (Supplementary Note 1). SALSA2 was found to be the best scaffolder and produced the closest agreement with the latest cattle reference ARS-UCD1.2. The scaffold N50 produced by SALSA2 was larger than those generated by optical map scaffolding, but the latter had the advantage of more accurate chimeric haplotigs detection, which resulted in 29 and 36 breaks in the Angus and Brahman haplotigs, respectively (Supplementary Note 2). Without these chimeric breaks, four apparent, incorrect inter-chromosomal fusions created in the initial haplotigs, involving two Brahman chromosomes (13 and 15) and six Angus chromosomes (8, 9, 12, 20, 23, 28), would have remained unresolved.

**Figure 1.**
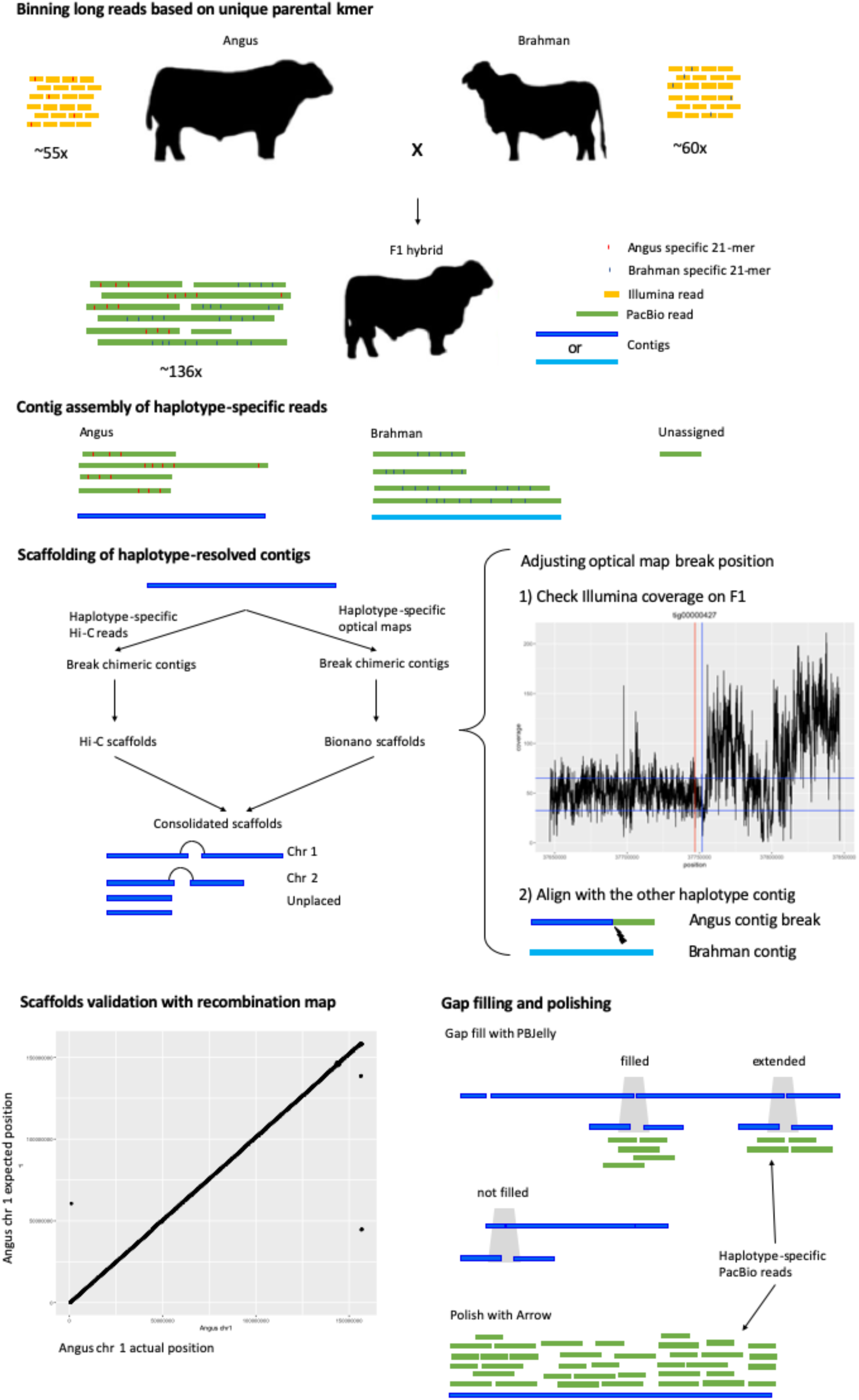
An overview of assembly methods. Long PacBio reads were binned to the respective haplotypes using parental specific k-mers prior to assembly by TrioCanu. Unassigned reads were discarded. Each haplotig assembly was scaffolded separately with Hi-C and optical map data. Both Hi-C and optical map scaffolding have chimeric haplotig detection capability and the optical map is better at resolving chimeras. Where possible the break positions of optical map-based cuts were adjusted based on F1 coverage and alignment with the other breed haplotig. Hi-C and optical map-based scaffolds were consolidated and cattle recombination maps were used to validate the assembly. Finally, haplotype specific long reads were used to fill gaps and polish the sequence.

After validation against a recombination map, gap filling, and error correction, the final assemblies, UOA_Angus_1 and UOA_Brahman_1 had similar chromosome sizes and excellent co-linear chromosome alignment with the current cattle reference, ARS-UCD1.2 (Supplementary Fig. 2). Unlike some of the recent PacBio-based assemblies^8, 9^, which required an additional polishing step with Illumina short reads to correct the high indel error rates, the haplotype-resolved assemblies only required correction of a very small number of coding sequences, showing that polishing with short reads was unnecessary (Supplementary Note 3).

The Brahman genome was annotated by Ensembl and NCBI whereas the Angus genome was annotated only by Ensembl (Supplementary Note 3-4). A comparison of annotation features between the Angus, Brahman and Hereford reference genomes is given in Supplementary Table 1. As the Ensembl pipeline was used to annotate all three cattle genomes, interpretation of results reported here used Ensembl release 96.

### Assembly benchmarking and sequence contiguity assessments

The per-base substitution quality values (QVs) for the UOA_Angus_1 and UOA_Brahman_1 reference assemblies were 44.63 and 46.38, respectively (Supplementary Table 2, Supplementary Note 5). The QV represents the phred-scaled probability of an incorrect base substitution in the assembly, hence these QVs indicate that the assemblies are more than 99.99% accurate at single base level. This is similar to the latest water buffalo assembly UOA_WB_1 (QV 41.96) and surpasses the recent goat ARS1 assembly (QV 34.5) by an order of magnitude. The Angus and Brahman assemblies had ∼93% BUSCO completeness score, which demonstrates a high-quality assembly of genes (Supplementary Table 3).

The Angus and Brahman assemblies have few gaps compared to most existing mammalian reference assemblies, and are comparable to the human GRCh38, the new Hereford cattle ARS-UCD1.2 and the water buffalo UOA_WB_1 reference genomes (Fig. 2a). For example, the Angus chromosome 24 was assembled without gaps. In terms of contiguity, these new cattle reference genomes are comparable to the recent water buffalo UOA_WB_1 assembly^9^, which is the most contiguous ruminant genome published to date with fewer than 1000 contigs (Supplementary Fig. 3), although it is not fully haplotype-resolved. While the cattle autosomes showed excellent contiguity, the Brahman X and Angus Y chromosomes were interrupted by 91 and 69 gaps, respectively.

**Figure 2.**
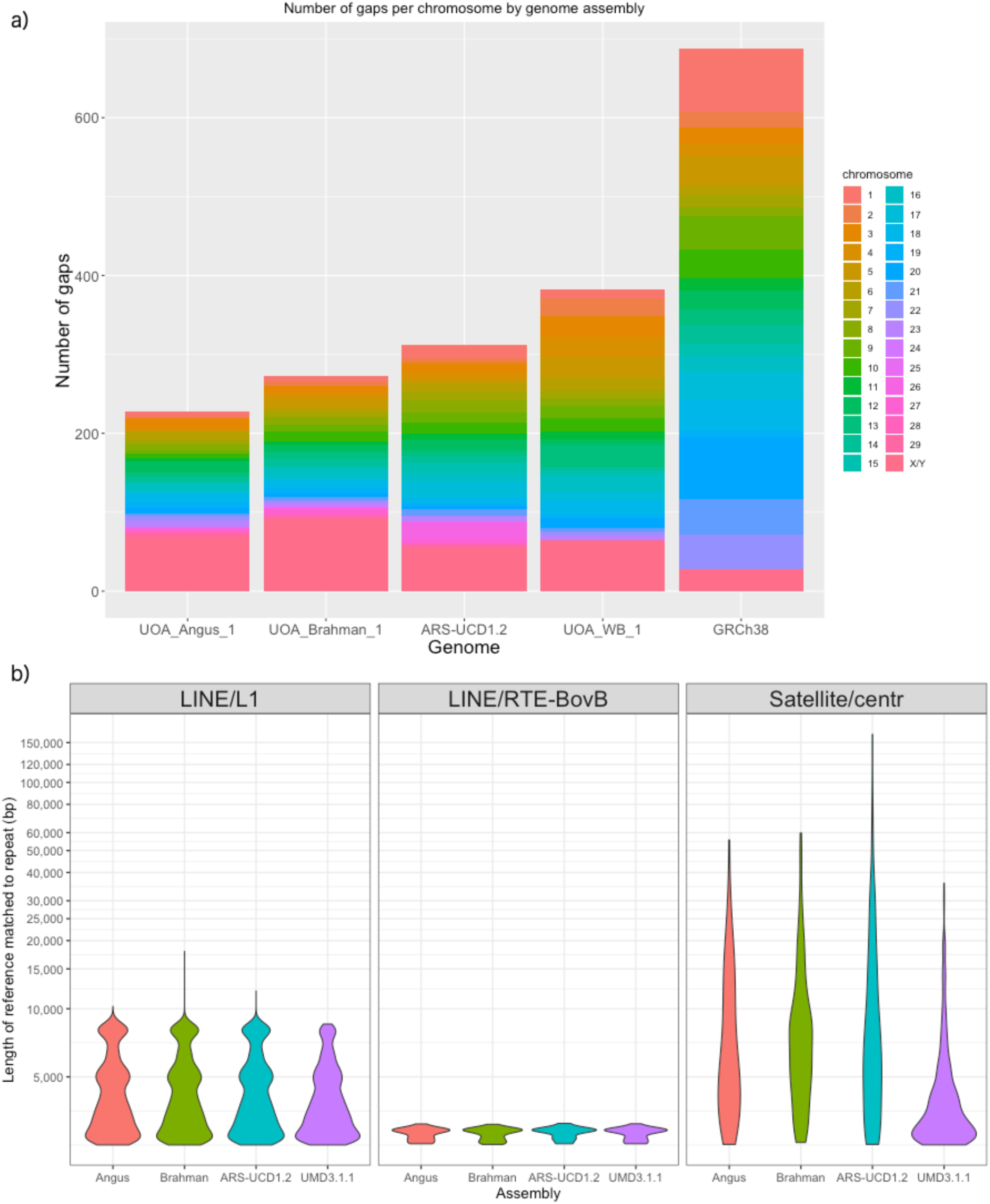
Sequence contiguity and resolution of repeats. a) Barplot of the number of gaps by chromosomes between various mammalian assemblies b) Violin plot of repeat families filtered for those more than 2.5 kb for LINE/L1, LINE/RTE-BovB and satellite/centromeric repeats.

### Resolution of longer repeats

The use of long PacBio reads substantially improved repeat resolution compared with the previous cattle assembly UMD3.1.1, which was assembled from Sanger sequences^10^ (Fig. 2b). Approximately 49% of both Angus and Brahman assemblies consist of repeat elements, which is consistent with other published mammalian assemblies, including human GRCh38, Hereford cattle ARS-UCD1.2, water buffalo UOA_WB_1 and goat ARS1. The two largest repeat families identified were Long Interspersed Nuclear Element (LINE) L1 and LINE/RTE-BovB, which covered ∼25% of the chromosomes in both cattle sub-species. Satellite or centromeric repeats (>10 kb) accounted for 21% and 14% of repeats in unplaced scaffolds of Angus and Brahman, respectively (Supplementary Fig. 4). The 7% higher satellite and centromeric repeats in Angus unplaced scaffolds is likely due to the presence of the Y chromosome in the Angus haplotype. The combination of the three most frequent repeat families, LINE L1, LINE/RTE-BovB and satellite/centromeric repeats, covered ∼40% of all unplaced bases, and repeat sequences were most frequently responsible for breaking sequence contiguity. The three cattle assemblies constructed using PacBio long reads that resolved repeats > 2.5 kb, UOA_Angus_1, UOA_Brahman_1 and ARS-UCD1.2, provide significant improvements in repeat resolution over the previous Sanger-based cattle assembly (UMD3.1.1) (Fig. 2b). All 29 cattle autosomes are acrocentric and in these assemblies, 20 contained centromeric repeats within 100 kb of chromosome ends, demonstrating that they approach chromosome-level. Nine Angus and eight Brahman chromosomes have centromeric repeats larger than 10 kb near the chromosome end. Six vertebrate telomeric repeats (TTAGGG)n were found within 1000 kb of chromosomal ends in Angus and five in Brahman assemblies.

### Discovery of indicus-specific fatty acid desaturase 2

One of the most diverged genomic regions between Brahman and Angus was observed on chromosome 15 (Fig. 3a). A region of ∼1.4 Mb has three copies of fatty acid desaturase 2-like genes (*FADS2P1*) in Brahman whereas the homologous region in the Angus only has two *FADS2P1* genes (Fig. 3b, c). In both Brahman and Angus the *FADS2P1* genes are encoded by 10 to 12 exons and the entire regions were assembled completely without gaps for both genomes. The region also contains six genes annotated as olfactory receptor-like, with unknown functions, which had differences in their predicted gene models between Brahman and Hereford assemblies. Within the ∼1.4 Mb region, there is a high level of sequence divergence for ∼200 kb, which is where an extra copy of *FADS2P1* lies in Brahman. Searches for *FADS2P1* in other ruminant species with high-quality genome assemblies revealed that only Brahman has three copies of the gene. The additional *FADS2P1* gene is ∼53 kb long, and is flanked by two other conserved *FADS2P1* genes. Searching WGS short read sequences from 38 animals used in this study showed that only Brahman animals had the extra copy, which was not present in any of the taurine individuals. (Supplementary Fig. 5). Considering that the Brahman genome is derived from four indicine breeds, the extra *FADS2P1* is likely a *Bos taurus indicus*-specific gene. We used CODEML to search for positively selected amino acid residues in FADS2P1 and identified 16 significant positively selected sites, 10 of which are located in a small exon 7 of only 60 bp (Fig. 3d, Supplementary Table 4).

**Figure 3.**
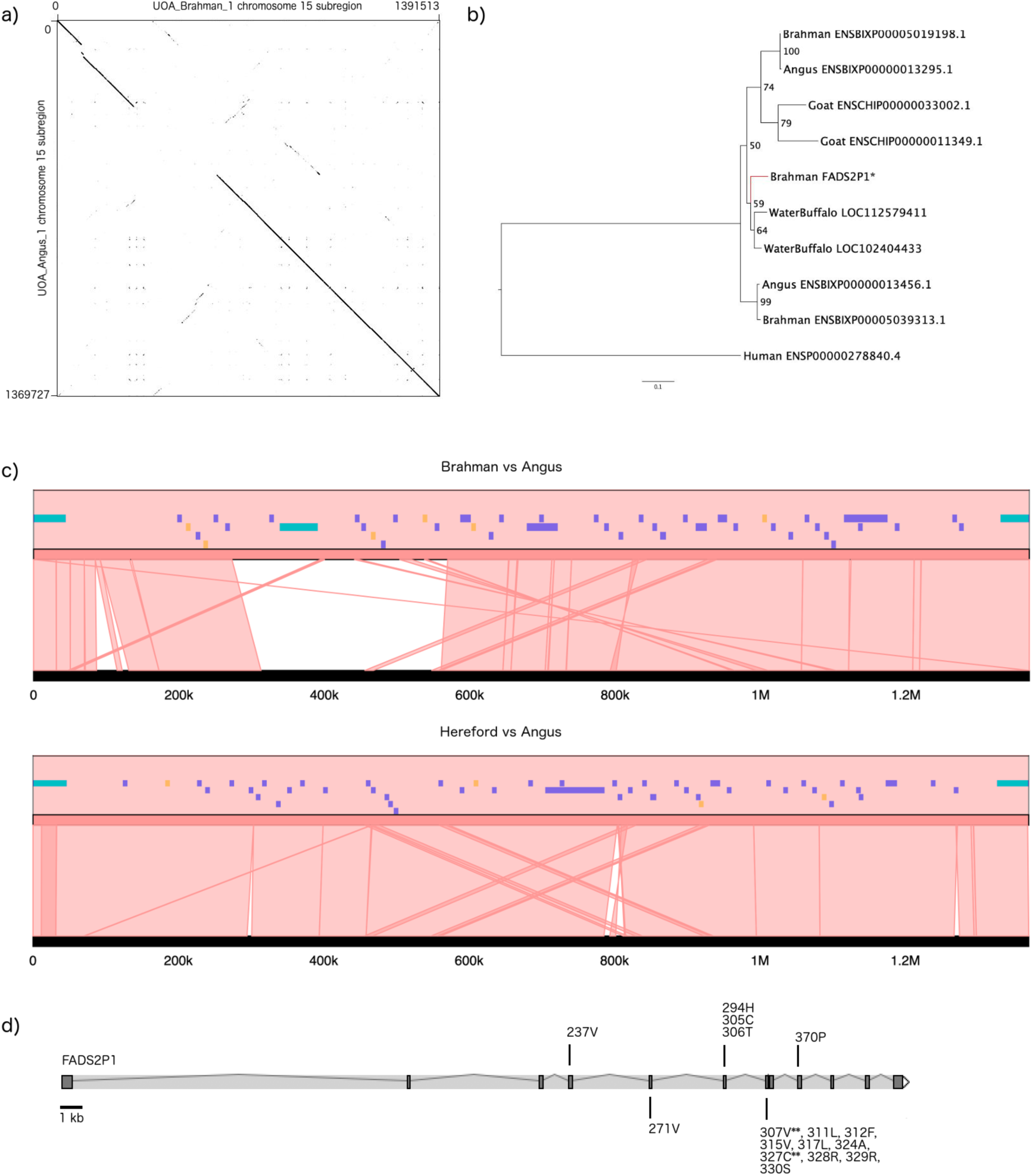
Discovery of *FADS2P1* locus as a divergent region between indicine and taurine cattle. a) Dot plot of Brahman chromosome 15 between positions 3,748,952 to 5,140,465 against the homologous Angus chromosome between positions 78,799,177 to 80,168,904. The Brahman sequence was reverse complemented in the plot. b) Maximum likelihood tree with 1000 bootstraps of FADS2P1 homologous protein sequences. The extra Brahman FADS2P1 copy is highlighted with “*” and its branch colored red. c) Microsynteny plot showing a lack of sequence conservation between indicine and taurine breeds at the indicine-specific *FADS2P1* gene. All *FADS2P1* genes are colored in turquoise, other genes in purple and pseudogenes are colored in orange. The upper plot used Brahman as the reference whereas the lower plot used Hereford as the reference. In each panel, the reference is always at the top. The Brahman *FADS2P1* genes Ensembl IDs are ENSBIXG00005007613, ENSBIXG00005021668 and ENSBIXG00005022680 whereas the Angus IDs are ENSBIXG00000018262 and ENSBIXG00000018381. d) Mapping of 16 positively selected sites onto the exons of Brahman *FADS2P1*. The residues with “**” indicate they have *prov*(ω = 1) > 0.99 (i.e. highly significant positively selected sites).

### Selective sweeps in Brahman

We compared the SNP patterns of 5 Brahman with 33 individuals from six taurine breeds to identify signatures of selective sweeps (Fig. 4a, Supplementary Note 6). We searched 100 kb windows spanning the whole genome for those where there was high level of homozygosity within Brahman individuals but with more segregating polymorphic variants in taurine breeds. This identified a total of 128 genes in 60 selective sweep intervals. Among these candidate selected genes, 80% were protein-coding, 1 was a pseudogene and the remainder were RNA-based genes (Supplementary Table 5). No biological pathways were found to be significantly over-represented among the positively selected protein-coding genes. The heat shock protein HSPA4, a member of the Hsp70 family, was amongst genes identified as under selection, which was also found in a search for selective sweeps in African cattle^11^. We also identified *DNAJC13*, a member of a gene family known to act as co-chaperones of heat-shock proteins, in another selective sweep region. In addition to heat tolerance related genes, selective sweep regions included genes involved in a range of biological processes including metabolic process, cellular component organization or biogenesis, cellular process, localization, reproduction, biological regulation, response to stimulus, developmental process, rhythmic process, biological adhesion and multicellular organismal process. Candidate genes in these regions under selection include two hormone receptors (*CRHR1, THRB*), two growth factor receptors (*FGFRL1, IGF1R*), immune related genes (*IL17B, IL10*) and an early growth response 2 (*EGR2*) transcription factor.

**Figure 4.**
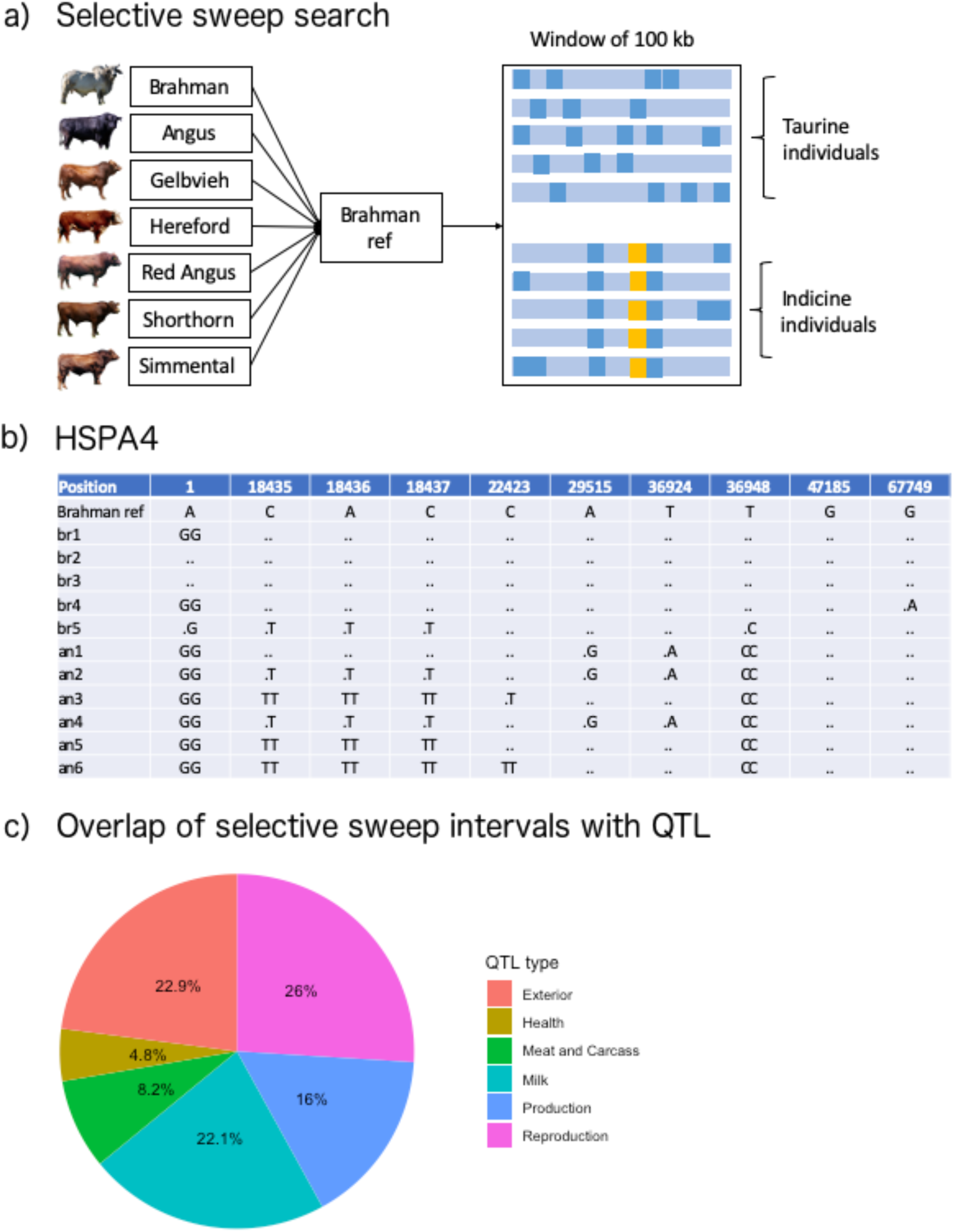
Selective sweep analysis in Brahman. a) An overview of the strategy used to identify selective sweep regions by aligning and calling SNPs in 100-kb windows on the Brahman reference. b) An example of genotype calls for *HSPA4*, a gene residing in one of the selective sweep intervals, using partial results from Angus and Brahman individuals. Genotype shown as “.” means it follows the Brahman reference, which is haploid, and “an” denotes Angus whereas “br” denotes Brahman. The position indicates adjusted position starting from 66,130,388 bp in UOA_Brahman_1, chromosome 7. c) Overlap of selective sweep intervals with cattle QTL categorized by types. Only unique QTL IDs were used.

A total of 231 unique QTL covering all six major QTL types listed in the cattle QTL database overlapped with positively selected genomic regions. The major QTL types were reproduction (26%) followed by exterior phenotype (22.9%) and milk (22.1%) traits (Fig. 4c). Ten of 60 selective sweep intervals did not overlap with any of the currently identified QTL.

### SNP and INDEL differences between Brahman and Angus

Using WGS short reads from 5 Brahman and 6 Angus individuals, we identified ∼24 million Brahman SNPs and ∼11 million Angus SNPs, which were annotated using the corresponding reference genomes (Table 2, Supplementary Table 6). There were about twice as many INDELs in the Brahman (2,804,421 bp) than the Angus (1,381,548 bp) genome. Mapping short reads from Brahman and Angus to both reference genomes, UOA_Brahman_1 and UOA_Angus_1, revealed that the use of breed-specific reference genomes gave a more accurate count of genetic variants. Lower false positives of SNPs, INDELs, and the four classes of structural variants (i.e. BND, DEL, DUP, INV) were identified when the appropriate reference genome was used. For example, a lower count of SNPs, by ∼4%, was observed when Brahman individuals were mapped onto the Brahman instead of the Angus reference genome.

**Table 2:** Polymorphism statistics

| Breed<sup>a</sup> | Reference | SNP | INDEL | BND | DEL | DUP | INV |
| --- | --- | --- | --- | --- | --- | --- | --- |
| Angus | Angus | 10615122 | 1381548 | 38 | 84 | 22 | 3 |
| Brahman | Angus | 24930357 | 2928526 | 311 | 641 | 86 | 22 |
| Angus | Brahman | 16504067 | 2090735 | 159 | 182 | 40 | 11 |
| Brahman | Brahman | 23876357 | 2804421 | 279 | 481 | 97 | 18 |

<sup>a</sup>Breed here refers to the ~10x WGS short reads used to align to the reference. BND, DEL, DUP and INV are structural variant types called in Lumpy. (BND = complex structural variant; DEL = deletion; DUP = duplication; INV = inversion).

### Structural variant differences between Brahman and Angus

We assessed the structural continuity of our Brahman and Angus genome assemblies against the current cattle reference genome assembly, ARS-UCD1.2, and against WGS datasets from 38 animals representing seven breeds, to ascertain the benefit of using haplotype-resolved assemblies for variant calling. To assess structural variant (SV) differences between Brahman and Angus and the cattle reference genomes, the haplotype-resolved assemblies were aligned to the ARS-UCD1.2 reference (Hereford). This detected insertions, deletions, tandem expansions, tandem contractions, repeat expansions and repeat contractions^12^ (Supplementary Fig. 6). Both tandem expansion/contraction and repeat expansion/contraction are repeat-type SVs. Detection of SV was limited to sizes of 50-10,000 bp, and the total bp affected by SVs in Angus and Brahman were 10.9 Mb and 21.8 Mb. This translates to approximately 0.4% and 0.8% of the Angus and Brahman genomes, respectively. Among the six classes of SVs examined, insertion/deletion types were the most prevalent in both Brahman and Angus genomes compared to ARS-UCD1.2.

We extracted Brahman- and Angus-specific SV to study their distribution in genic and intergenic regions (Fig. 5a). For Brahman-specific SV, insertion/deletion types overlapped ∼4% of all genes whereas each of the repeat-type SV overlapped ∼1% of genes. In contrast, the Angus-specific insertion/deletion type SV overlapped ∼1-2% of all genes and the repeat-type SV overlapped less than 1% of genes. Therefore, the majority of SV were found in intergenic regions and whenever they overlapped with genes were generally localized within introns. Over-representation of Gene Ontology (GO) terms were detected for Angus-specific insertions and tandem contractions and Brahman-specific insertion/deletion SVs at FDR-adjusted P-value < 0.05 (Supplementary Table 7). No over-representation of GO terms was detected for any of the other breed-specific SV types. Interestingly Brahman-specific insertion SVs have between 3 to 5.7 fold enrichment in phospholipid translocation (GO:0045332), lipid translocation (GO:0034204), lipid transport (GO:0006869), and lipid localization (GO:0010876) GO classes, which suggests lipid distribution was most impacted by SVs.

**Figure 5.**
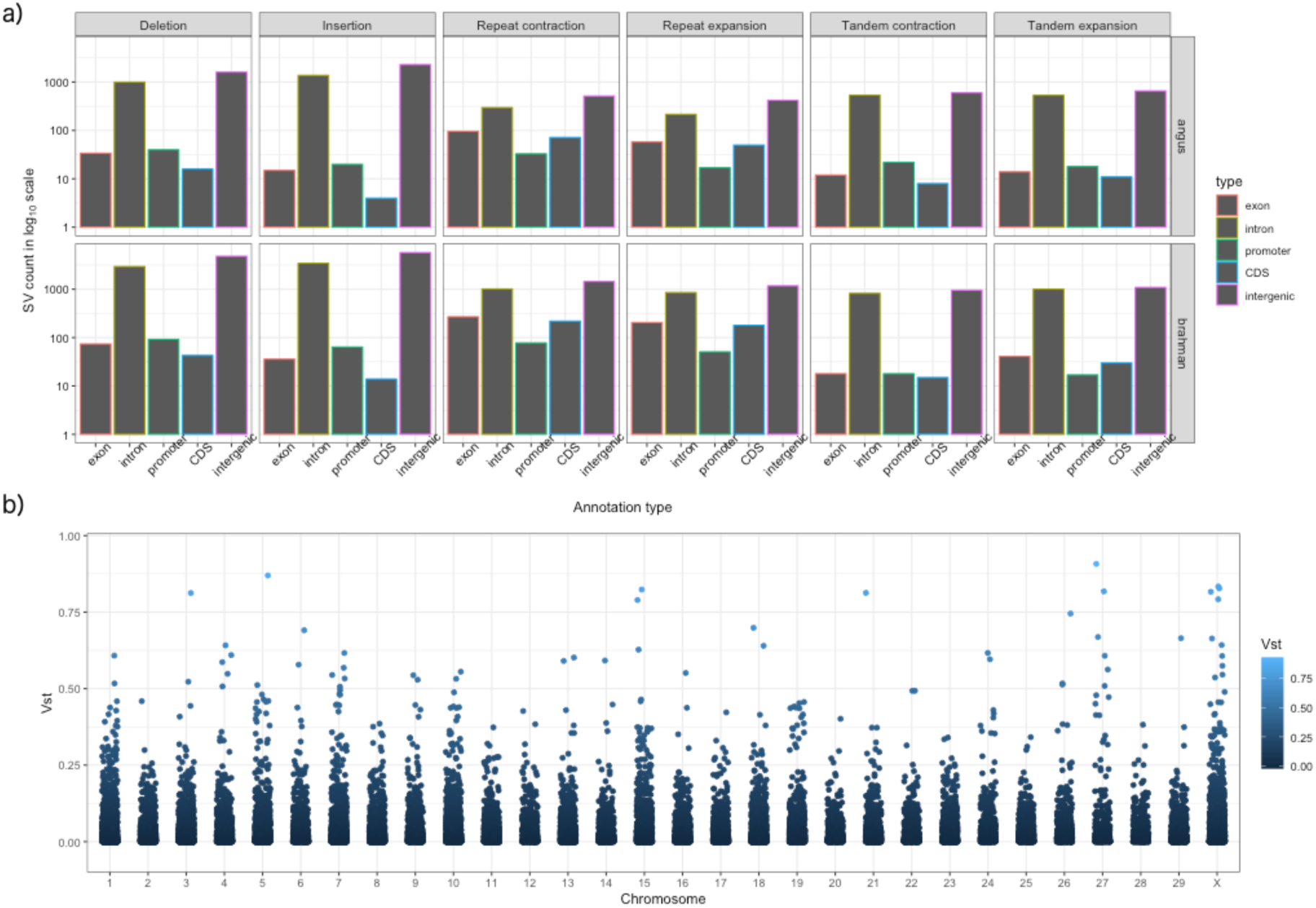
Comparison of structural variants between Brahman and Angus. a) Count in log_10_ scale of 6 classes of SVs when overlapped with various annotation types. b) Population differentiation for copy number variation as estimated by *V_ST_* along each chromosome for the taurine and indicine comparison using UOA_Brahman_1 as the reference.

Using WGS reads from different datasets, we used a two-tiered approach to identify subspecies-specific CNVs that were masked by the absence or poorer resolution of sequence in the ARS-UCD1.2 reference. The input dataset for these analyses came from ∼10x WGS short reads of 38 animals representing seven cattle breeds. Each set of reads was aligned to all three reference genome assemblies (Hereford, Brahman, and Angus) and processed with SV callers designed to detect read depth differences and paired-end/split-read discordancy, respectively. The first approach (read-depth variation) included the use of the V_st_ statistic^13, 14^ to identify genes with copy number variation between taurine or indicine lineages using the Brahman, Angus or ARS-UCD1.2 assemblies, (Fig. 5b, Supplementary Fig. 7). Only autosomes were considered. Six CNV genes were found in Brahman whereas four and eight CNV genes were found in Angus and Hereford, respectively (Fig 6a-c). Prediction of CNV genes was sensitive to the assembly chosen, e.g. only *TMPRSS11D* and beta-defensin-like precursor were found to be copy number variable in more than one assembly. Among the 18 CNV genes differentiating indicine from taurine genomes, six unique gene families were identified, which were beta defensin, workshop cluster, trypsin-like serine protease, T-cell receptor alpha chain, tachykinin receptor, and interferon-induced very large GTPase, all of which have immune-related functions. All of the CNV genes from these six families showed higher copy number in the indicine cattle lineage regardless of the assembly used. An olfactory receptor, two long non-coding RNAs and one putative protein, FAM90A12P, also had higher copy numbers among indicine animals. In contrast, ubiquitin-conjugating enzyme E2D3 and two keratin-associated protein 9 genes (*KRTAP9-1, KRTAP9-2*) had higher copy numbers in the taurine lineage.

**Figure 6.**
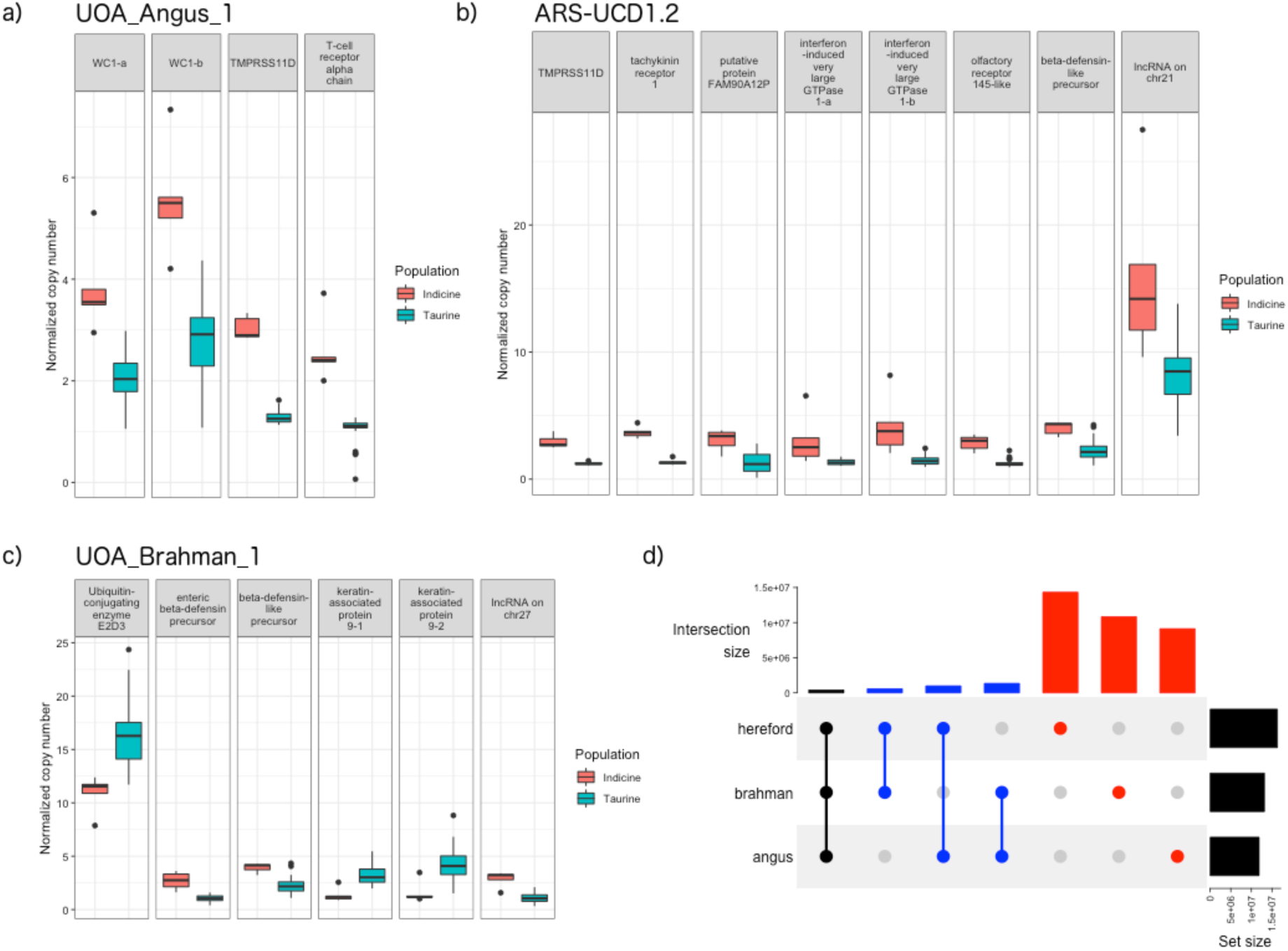
Boxplot of normalized copy number of autosomal genes with *V_ST_* greater than 0.3. Only those CNV genes with average copy number difference of at least 1.5 copies between taurine and indicine groups are shown. The reference genomes were a) UOA_Angus_1, b) ARS-UCD1.2, and c) UOA_Brahman_1. d). Liftover of CNV regions from Brahman and Angus to Hereford ARS-UCD1.2 common coordinate for an assessment of intersection between them at base-pair resolution.

We quantified the effects of using different reference assemblies for paired-end/split-read (PE) SV discovery. All SV calls of this type were converted into Hereford coordinates to facilitate comparisons. We removed 17, 9, and 18 PE SVs from the Brahman, Angus and Hereford assemblies that were likely false positives, as they were larger than 1 Mb and did not correspond to aberrant read depth signal to support their SV calls. On average, 0.5% of each cattle genome was covered by CNV regions (CNVRs) (Fig. 6d). The majority of CNVRs (at least 76% from each assembly) were found to be unique to one assembly. Among the Brahman CNVRs, only 10% intersected with Angus CNVRs, which suggests mis-assembly in the Hereford reference potentially due to compression of repetitive elements that are more difficult to resolve without phasing haplotypes using the trio binning method.

### Phasing of full-length transcripts in haplotype-resolved genomes

Among the PacBio error corrected Iso-Seq (CCS) reads pooled from seven tissues of the F1 hybrid fetus, 3,275,676 reads (55%) were classified as full-length non-concatamer (FLNC) reads. After processing with the isoseq3 software, 193,974 full-length, high-quality (HQ) consensus transcripts were generated. We mapped the HQ transcripts to the Brahman reference and obtained 99,329 uniquely mapped transcripts covering 20,940 non-overlapping loci. Using the SQANTI2 transcript characterization tool, 83% of the Iso-Seq transcripts fell into coding regions of the Brahman annotation (Fig. 7a). As many as 68% of the transcripts could be considered as novel, because they were categorized as “novel in catalog” (NIC), “novel not in catalog” (NNC), antisense, intergenic or genic. The transcript length distribution ranged from 85 to 11,872 bp, with a median of 3853 bp and a mode of ∼4 kb (Fig. 7b).

**Figure 7.**
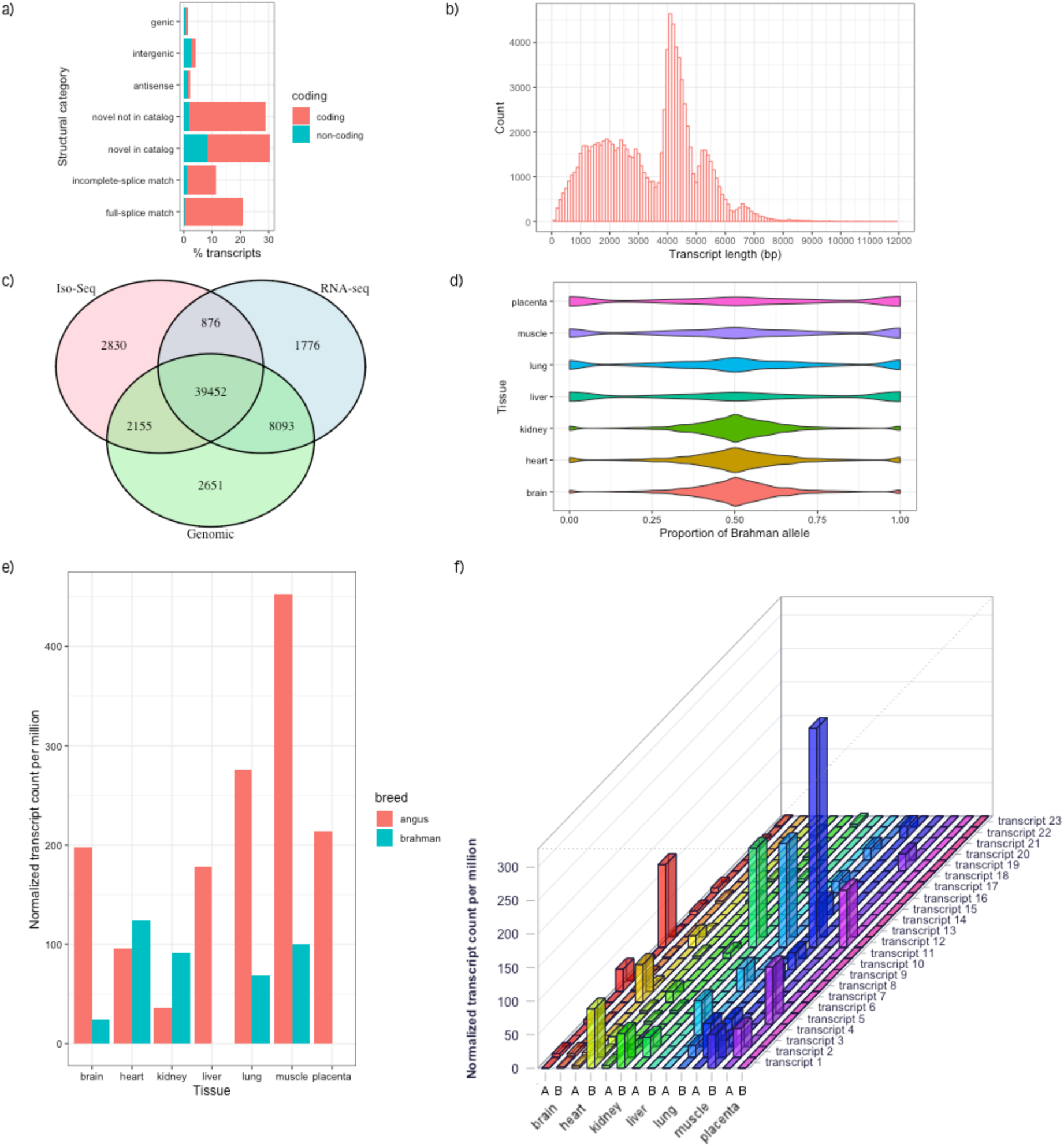
Phasing of Iso-Seq full-length transcripts in seven tissues reveals transcriptional complexity and allelic imbalance. a) Characterization of transcript annotation of the hybrid animal using SQANTI2 against the Brahman annotation. Full-splice match: perfect match with a reference; Incomplete-splice match: missing one or more 5’ exons against a reference; novel in catalog: novel combinations of known junctions; novel not in catalog: at least one novel splice site. b) Histogram of transcript length distribution. c) The overlap of SNPs between WGS short reads from genomic DNA, Iso-Seq and RNA-Seq when Brahman was used as the reference genome. d) Violin plot of the proportion of Brahman alleles, which was calculated as the normalized count of Brahman alleles divided by the sum of normalized count of both Brahman and Angus alleles. Transcripts showing allelic imbalance and with higher expression in Brahman have values closer to 1 whereas those with higher expression in Angus have values closer to 0. e) Tissue-specific allelic expression at the gene level for *ARIH2*, which is the most highly expressed Angus gene in the brain f) Tissue-specific allelic expression at the transcript level for *ARIH2* in brain, heart, kidney, liver, lung, muscle and placenta. A denotes “Angus” and B denotes “Brahman”.

After exclusion of SNPs with less than 40-fold Iso-Seq read coverage and those in non-transcribed regions, IsoPhase identified 5806 genes with 52,270 phased transcripts. Of the 45,313 SNPs called by IsoPhase, 39,452 (87%) were validated by SNPs from RNA-Seq and genomic DNA, whereas 876 (1.9%) were only validated by RNA-Seq data and 2155 (4.7%) were only validated by genomic sequence data (Fig. 7c) (Supplementary Note 7). SNP calls that showed inconsistencies could often be explained by lower Iso-Seq coverage, SNPs in homopolymer regions, or alignment artefacts.

Our haplotype-resolved genomes allowed us to explore genes with allelic imbalance in expression. (Fig. 7d). All tissues showed evidence of imbalance in allelic expression (Shapiro test, P-value < 0.01), which was most pronounced for liver, lung, muscle and placenta, whereas brain, heart and kidney were less affected. However, as mammalian brain consists of a wide range of cell types and hence transcriptional complexity, brain tissue was chosen to demonstrate the phasing of transcripts to explore allele-specific expression. The most highly expressed Angus gene with allelic imbalance (ratio of 8 Angus : 1 Brahman) in the brain was *ARIH2* (also known as *TRIAD1*), which is known to play a role in protein degradation via Cullin-RING E3 ubiquitin ligases^15^ (Fig. 7e,f). *ARIH2* expression in the liver, lung, muscle and placenta was also higher from the Angus allele than the Brahman or maternal allele. The HQ transcripts included 23 different transcript isoforms of *ARIH2*, however, 66% of transcripts for this gene across the seven tissues were represented by only three isoforms. The annotated exons of this gene were in good agreement with the RNA-Seq data (Supplementary Fig. 8).

The most highly expressed Brahman gene with allelic imbalance (ratio of 1 Angus : 6 Brahman) in the brain was Calmodulin (*CaM*), a heat-stable Ca^2+^-binding protein that mediates the control of numerous physiological processes, including metabolic homeostasis, phospholipid turnover, ion transport, osmotic control, and apoptosis^16^ (Supplementary Fig. 9). Surprisingly, we also found allelic imbalance (ratio of 1 Angus : 16.5 Brahman) in pregnancy-associated glycoprotein 1 (*PAG1*) with a higher expression of the Brahman allele in the brain and placenta but undetectable in other tissues. This gene was previously thought to be placenta-specific and is used as a biomarker for embryo survival^17^.

## Discussion

Traditional genome assembly approaches collapse haplotypes and therefore do not allow accurate assembly or the study of divergent, heterozygous regions. Here we demonstrate a new assembly approach that yielded highly contiguous, haplotype-resolved Brahman and Angus cattle genomes from an F_1_ hybrid of the two subspecies. Our analyses demonstrated that previous studies, which mapped indicine sequences onto the taurine reference UMD3.1.1^4, 11^, likely inflated the number of genetic variants that are present between the two subspecies by up to 4%. Calling SNPs in transcripts from a diploid hybrid with both haplo-genomes decoded provides accurately phased transcripts for studies on the role of allele-specific expression in, e.g., hybrid vigor or heterosis. The phasing of Iso-Seq transcripts in reciprocal crosses will facilitate the exploration of breed-specific effects on parental imprinting, which has been shown in maize^18^.

We found that the choice of reference assembly had a large impact on SV calling. After converting SVs from each assembly onto the Hereford assembly coordinates and calculating the intersection, we identified 1.3 Mbp SVs present in the Angus and Brahman assemblies were not present in the Hereford assembly. This suggests that either the Hereford assembly was not as representative of the true structural variation in these regions or that there were assembly errors in the Angus and Brahman assembly that generated false positive SVs. The latter is less likely given the high accuracy of the Angus and Brahman genomes. Conversely, we identified 0.9 Mbp SVs shared between only the Hereford and Angus assembly, which may represent true genomic structural differences between taurine and indicine cattle.

The discovery of an indicus-specific, additional copy of fatty-acid desaturase 2 gene (*FADS2P1*), that has been under positive selection, further highlights the benefits of high-quality haplotype-specific assemblies. The *FADS2P1* gene region in both Brahman and Angus span ∼1.4 Mb of sequence, while the two *FADS2P1* genes in the water buffalo span ∼1 Mb. The orthologous region in goat is ∼1 Mb, but contains gaps. Taking phylogenetic and information on conservation of synteny together, the most parsimonious explanation is that the extra *FADS2P1* was duplicated in the indicine lineage after divergence from taurine cattle. Rapid evolution at the *FADS2P1* locus resulted in neofunctionalization of the additional gene in indicine animals, with profound changes seen in the small exon 7.

*FADS2* is a pleiotropic gene with known functions in the biosynthesis of unsaturated fatty acids, lipid homeostasis, inflammatory response, and promotion of myocyte growth and cell signaling^19–21^. A non-synonymous SNP in exon 7 of Japanese Black cattle is significantly associated with linoleic acid^22^ composition. While we do not know the functional significance of positively selected residues in the additional *FADS2P1* copy in Brahman, the SNP reported in the Japanese Black shows the importance of exon 7 in FADS2 function. Studies in rats have shown linoleic acid is an important component of skin ceramides and its deficiency increases water permeability of the skin^23^. Comparisons between indicine and taurine animals have shown differences in fatty acids^24^ and types of phosphatidylcholines^25^. We hypothesize *Bos indicus* has three copies of *FADS2P1* genes to regulate the composition of fatty acids that constitute the cell membranes and could alter water permeability and heat loss from skin.

The significant differences in phenotype, energy metabolism and adaptation to heat stress of indicine cattle have been linked to the thyroid hormone axis^26–28^. One of the selective sweep regions in Brahman contains thyroid hormone receptor β (THRB), a ligand-activated pleiotropic transcription factor that modulates the expression of a large number of genes^29^. Thyroid hormones are intrinsically connected to the growth hormone - insulin-like growth factor axis (GH-IGF)^30^. Insulin-like growth factor 1 receptor (IGF1R) was found in another selective sweep region. Polymorphisms in *IGF1R* have been associated with age of puberty in Brahman cattle^31^. In comparison with taurine cattle Brahman tend to reach puberty late, which may have been under positive selection as a consequence of adaption to harsh tropical environments, ensuring that cows are more mature and robust at the time of first calving.

Quantitative trait loci for reproduction featured prominently in the comparison of Brahman selective sweep regions with known cattle QTL. Amongst the reproduction traits, QTL related to calf size and calving ease were overrepresented. Brahman cows deliver a small calf that is less likely to result in dystocia and still birth^32^, which is one major benefit of the introgression of indicine genetics into more productive taurine breeds. Selective sweep regions thus provide candidate genes for maternal control of birth weight.

Brahman cattle may be better adapted to harsher environments because they have slower protein turnover^33^. Relative to Angus, Brahman have much lower expression of *ARIH2* in key metabolic organs, such as the skeletal muscle and no detectable expression in the liver. ARIH2 promotes ubiquitylation of DCNL1, which is a co-E3 ligase that performs cullin neddylation, a process that regulates one-fifth of ubiquitin-dependent protein turnover^15^. CNV analysis revealed a decreased number of ubiquitin-conjugating enzyme E2D3 genes in the indicine lineage, which suggests lower protein turnover in indicine animals. As a result, Brahman possibly have lower endogenous energy expenditure and protein turnover, and thus are better able to withstand stressful conditions than taurine cattle.

The CNV analyses of Brahman and Angus genomes revealed that six gene families with immune related functions and putative roles in response to disease challenge and external parasites are expanded in the indicine lineage. Conversely, *KRTAP9-*2, a gene with significantly altered gene expression following tick infestation^34^, is expanded in the taurine lineage, which has also been reported in previous CNV studies^14, 35^. Further studies are needed to elucidate how changes in copy number of *KRTAP9-*2 affect its expression and its role in tick resistance.

In conclusion, the approach used here is able to create haplotype-resolved genome assemblies that are of higher quality than traditional haplotype-collapsed assemblies. Availability of these high-quality assemblies has enabled us to better resolve structural variants and identify regions under selection that may be involved in adaption to the environment. Looking forward, it is clear that high-quality haplotype-resolved assemblies together with long-read transcripts information will underpin studies on genome function, regulation and the control of phenotypes.

## Methods

### Bos taurus hybrid

A *Bos taurus indicus* female (Brahman) was inseminated with semen from a *Bos taurus taurus* (Angus) bull. The *indicus* maternal genetic background of the Brahman dam was confirmed by mitochondrial DNA haplotype analysis^36^. At day 153 post-insemination, dam and conceptus were ethically sacrificed and fetal brain, heart muscle, kidney, liver, lung, skeletal muscle and placenta (cotyledon) tissue were snap frozen in liquid nitrogen and stored at −80°C until further use. All animal work was approved by the Animal Ethics Committee of the University of Adelaide (No. S-094-2005).

### Genome sequencing and assembly of contigs

DNA was extracted from fetal lung, dam uterus and bull semen as described previously^6^. Twelve SMRT sequencing libraries were made from the fetal DNA using the protocol recommended by the Pacific Biosciences (Procedure P/N 100-286-000-07), with a 15 kb size selection cut-off on a Blue Pippin instrument (Sage Science, Beverley, MA). Nine libraries were sequenced using P6/C4 chemistry on an RSII machine whereas the remaining three libraries were sequenced on a Sequel machine. Approximately 161 Gb of RSII data and 205 Gb of Sequel data were produced, which gave a total sequence yield of 366 Gb with the mean read length of ∼10.4 kb. Assuming a genome size of 2.7 Gb, the raw PacBio data represents ∼136x coverage.

Illumina sequencing libraries for both parents (i.e. sire and dam) and F1 fetus were prepared using TruSeq PCR-free preparation kits (Illumina, San Diego, CA). A total of ∼55x, ∼60x and ∼84x coverage of 150 bp paired-end reads were generated for the sire, dam and F1 fetus, respectively. In order to assemble phased haplotigs for the F1 Brahman-Angus hybrid, we used the Trio-binning method introduced by Koren et al^6^. Briefly, 21-mers in both sire and dam Illumina reads were identified and 21-mers unique to one or other parent were used to assign the F1 PacBio long reads to the parent of origin. Approximately 1% of the PacBio reads were excluded from the assembly as they lacked parent-of-origin-specific 21-mers, due to their shorter lengths. Long reads that were binned into paternal and maternal groups were assembled separately with TrioCanu v1.6.

### Hi-C library preparation and sequencing

A Sau3AI Hi-C library was prepared (Phase Genomics, Seattle WA) as follows: approximately 200 mg of fetal lung tissue was finely chopped and then cross-linked in Proximo crosslinking solution. The 5’ overhangs after Sau3AI digestion were filled with biotinylated nucleotides, and free blunt ends ligated. After ligation, crosslinks were reversed and the free DNA was column purified and sonicated to approximately 600 bp peak fragment size (Bioruptor, Diagenode). Hi-C junctions were bound to streptavidin beads and washed to remove unbound DNA. Washed beads were used to prepare sequencing libraries using the HyperPrep kit (Kapa) following manufacturer’s protocols. In total, 203 million 2x 81 bp read pairs were sequenced on NextSeq Illumina platform.

### Scaffolding of contigs with Hi-C

All Hi-C reads were mapped to each breed-specific set of haplotigs using BWA^37^. A haplotype score for a pair was defined as the sum of the percent identity multiplied by match length for each read end (unmapped read ends were assigned a score of 0). Each read pair had two scores, one per haplotype. Pairs with a higher score for one haplotype were considered breed-specific and assigned to their respective haplotype. Pairs with a tied score were considered homozygous and assigned to both haplotypes for scaffolding.

Three different Hi-C based scaffolding programs, 3D-DNA^38^, Proximo (Phase Genomics) and SALSA2^39^ were evaluated for scaffolding contigs. Further detail on the comparison between the scaffolders is given in Supplementary Note 1. Reads were mapped with the Arima mapping pipeline (https://github.com/ArimaGenomics/mapping_pipelinecommit72c81901c671203a86ca4675457004a71d0cd249) and converted to bed format prior to SALSA2 scaffolding (https://github.com/machinegun/SALSAgitcommit863203dd094aaf9b342c35feedde7dabeec37b44), which was run with parameters ‘-c 10000 -e GATC -m yes’ for both breed-specific haplotigs.

### Bionano DNA isolation and assembly

DNA was extracted from 10 mg kidney tissue from the F1 hybrid using the Bionano Animal Tissue DNA Isolation Kit (P/N 80002) with slight modifications as follows: the frozen tissue was crushed in liquid nitrogen, placed in 2% formaldehyde in Bionano animal tissue homogenization buffer (Document number 30077, Bionano-Prep-Animal-Tissue-DNA-Isolation-Soft-Tissue-Protocol.pdf), and blended with a rotor-stator. The homogenate was passed through a 100 um nylon filter, fixed on ice for 30 minutes in 2 mL 100% ethanol, and centrifuged for 5 minutes at 2000 g. The resulting pellet was re-suspended in homogenization buffer and added to pre-warmed agarose to make 0.8% agarose plugs. High molecular weight DNA was extracted from the agarose plugs, labelled, stained, and imaged on a Bionano Saphyr system^40^. Further detail on *de novo* optical map assembly is given in Supplementary Note 2.

### RNA-Seq and Iso-Seq

RNA was extracted from tissue and ground to a fine powder under liquid nitrogen using the Qiagen RNeasy Plus Universal kit as per the manufacturer’s instructions. RNA quality was assessed using an Agilent TapeStation system and confirmed as RIN >8 for all samples. Sequencing libraries were prepared with the KAPA Stranded RNA-Seq Library Preparation Kit as per the manufacturer’s protocol and sequenced on an Illumina Next-Seq machine for 100 bp paired-end reads with the target of 50 million reads per sample.

Iso-Seq data were generated from brain, heart muscle, kidney, liver, lung, skeletal muscle and placenta (cotyledon) tissue. Iso-Seq SMRT bell libraries were created according to the PacBio protocols. Briefly, two size selected cDNA pools were created, one with an average cDNA size ∼ 3 kb and the second with a cDNA size of ∼7 kb. The two pools were then combined for SMRTbell™ Template Preparation. The final average library size was ∼5 kb as measured by a bioanalyser. Each SMRTbell library was loaded onto the Sequel at approximately 50pM.

### Identification and phasing of full-length transcripts

The Iso-Seq data was processed using the isoseq3.1.0 software on the PacBio Bioconda (https://github.com/PacificBiosciences/pbbioconda). The process consists of: (1) generating circular consensus sequence (CCS) reads, (2) classifying full-length non-concatamer (FLNC) reads that have the 5’, 3’ cDNA primer sequence and the polyA tail, (3) clustering FLNC reads at the isoform-level and generating a draft consensus for each isoform, and (4) polishing each isoform to create high-quality, full-length transcript sequences.

The high-quality transcript sequences were then mapped to the Brahman reference genome using minimap2 (v2.15-r905) and filtered for alignments that had ≥99% coverage and ≥95% identity. Redundant and degraded transcripts were collapsed using the Cupcake tool (https://github.com/Magdoll/cDNA_Cupcake). SQANTI2^41^ was used to annotate transcripts for various features such as known isoforms with full-splice match (FSM) or incomplete-splice match (ISM), novel isoforms in catalog (NIC) or not in catalog (NNC), and other novel genes that are antisense, overlap with intergenic or genic regions.

In order to phase transcripts using the Iso-Seq data, we ran IsoPhase, which is a part of the Cupcake tool, against the Brahman reference. IsoPhase first piles up the FLNC reads of all the isoforms of a gene and calls substitution SNPs using a one-sided Fisher exact test with Bonferroni correction at a P-value cut-off of 0.01. It then infers haplotypes based on the phasing information provided by the FLNC reads. The output defines the inferred haplotypes for each transcript and estimates the relative abundance of each allele. We ran IsoPhase using the pooled set of all FLNC reads from all tissues, then later demultiplex them to create an abundance matrix that is specific for each haplotype, per isoform FLNC count for each tissue. To compare the abundance of transcripts across tissues, we normalized the counts by dividing the FLNC counts for each haplotype-isoform by the total number of FLNC counts in that tissue, multiplied by a million to obtain the transcript per million (TPM) number.

IsoPhase was able to identify at least one SNP in 6273 of the genes. We then validated the IsoPhase SNPs using (1) SNPs called from RNA-Seq data of brain, liver, lung, muscle, placenta, and (2) Angus SNPs derived from mapping Illumina WGS short reads of the F1 hybrid to the Brahman reference. For RNA-Seq, read mapping was performed with Hisat2^42^ whereas the genomic short reads were mapped using BWA v0.7.15^37^. SNPs were called using GATK v4^43^. As the RNA-Seq had greater coverage than Iso-Seq and the SNPs called from genomic DNA included non-transcribed regions, only SNPs that were in positions covered by at least 40 full-length Iso-Seq reads were retained. The proportion of an allele from each breed, to assess allelic imbalance, was calculated as the normalized count of the Brahman allele divided by the sum of normalized count of both Brahman and Angus alleles.

### Scaffold validation with recombination map

Scaffold contiguity was assessed using a previously published recombination map^44^. Briefly, the recombination map probe sequences were aligned using BWA MEM to the scaffolds and the coordinates were arranged in a directed acyclic graph, using a custom script. A contiguity break between consecutive recombination map-ordered probes in the scaffolds was considered an error, however, we tolerated one mismatched probe in a window of three consecutive probes (Hamming distance = 1) to avoid false positive detection due to mapping ambiguity. Despite having Hi-C sequences, some scaffolds that belonged to chromosomes could not be joined together, which necessitated the use of recombination map markers to join and orientate these scaffolds.

### Gap filling and polishing

After checking scaffolds with recombination maps^44^, the Angus and Brahman scaffolds that contained 343 and 369 gaps, respectively, were gap filled with PBJelly^45^ v15.8.24 using haplotype-specific PacBio subreads. The default parameters of PBJelly were used, except for the support module, where the options “captureOnly and spanOnly” were used. This step closed 52 and 61 gaps in Angus and Brahman scaffolds, respectively. Two rounds of ArrowGrid (see URLs) was run to polish the scaffolds to give quality scores.

### Assembly evaluation and genome annotation

The assemblies were evaluated with BUSCO v2.0.1^46^ and other metrics that include compression/expansion (CE) errors. Annotations were created using the Ensembl gene annotation system^47^ and the NCBI pipeline. Further detail on the annotation process is given in Supplementary Notes 3-4 and for assembly evaluation, detail is given in Supplementary Note 5.

### Repeat analysis

RepeatMasker version open-4.0.7 (see URLs) was used to search for repeats in the UOA_Angus_1 and UOA_Brahman_1 assemblies by identifying matches to RepBase (version RepBase23.10.embl)^48^. Repeats in the current water buffalo assembly (UOA_WB_1) and cattle assembly (UMD3.1.1) were downloaded from the NCBI. Repeats with matches less than or equal 60% identity were filtered out. Centromeric repeats were identified by searching repeats that belonged to the family ‘Satellite/centr’ in Repbase. The vertebrate telomeric repeat, 6-mer TTAGGG, was identified by RepeatMasker. The search for at least 2 consecutive identical TTAGGG repeats within 1000 kb of chromosome ends was done to detect presence of telomeres.

### Gap comparisons and sequence contiguity

To evaluate gaps and sequence contiguity, the Angus and Brahman assemblies were compared to the water buffalo, human and Hereford cattle assemblies. Only sequences that belong to autosomes and sex chromosomes were retained for analysis, whereas unplaced and mitochondrial sequences were filtered out. The tool seqtk v1.2-r94 (see URLs) was used to count gaps with similar code implementation as those used for the water buffalo genome^9^.

### Single-nucleotide polymorphism and indel calls

38 individuals with ∼10x WGS short read Illumina data representing seven breeds were selected from the USMARC Beef Diversity Panel version 2.9 (MBCDPv2.9)^49^. The individuals selected for the panel were bulls with minimal pedigree relationships to maximize sampling of diverse alleles suitable for population genetics studies. The number of individuals per breed was as follow: six Angus, five Brahman, six Gelbvieh, six Hereford, five Red Angus, five Shorthorn and five Simmental. These six taurine breeds were chosen on the basis that they were unlikely to carry *Bos indicus* genetics given their history.

WGS data quality of each individual was checked with FASTQC v0.11.4^50^ and then trimmed with Trim Galore v0.4.2^51^ to a minimum length of 110 bp per read and Phred score of 20. Potential adapters in the sequence reads were removed using AdapterRemoval v2.2.1^52^. Following trimming, the reads were checked with FASTQC again to ensure that only high-quality reads were retained. Reads were then mapped to both the Angus and Brahman assemblies separately using BWA v0.7.15^37^ with the option “mem”. Samtools v1.8^53^ was used to convert the resulting alignment to sorted bam format. Duplicate reads, that may be due to PCR artifacts, were marked with Picard^54^ MarkDuplicates. The bam files from each individual animal were merged with GATK v4^43^ MergeSamFiles function. Then the following series of GATK functions, AddOrReplaceReadGroups, HaplotypeCaller, CombineGVCFs and GenotypeGVCFs, were applied to the alignment files to generate a variant call file in VCF v4.2 format. SNPs were filtered with VariantFiltration function using the parameters “(QD < 2.0) || (FS > 60.0) || (MQ < 40.0) || (MQRankSum < −12.5) || (ReadPosRankSum < −8.0)”. Indels were filtered with VariantFiltration function using the parameters “(QD < 2.0) || (FS > 200.0) || (ReadPosRankSum < −20.0)”. Annovar tool^55^ version dated 2017-07-17 was used to annotate the variants.

### Structural variant and copy number variant analyses

WGS short read data sets from the same 38 animals used for SNP and indel calls were aligned to the UOA_Angus_1, UOA_Brahman_1 and ARS-UCD1.2 with BWA MEM^37^ and further processed with Samtools v1.9^53^. Read-pair and split-read profile structural variants were called with the lumpy-sv v0.2.13^56^ pipeline, lumpyexpress, using default parameters for each sample. lumpy-sv VCF files were converted to BEDPE format using the vcfToBedpe script included in the lumpy-sv software package. Copy number estimates for genomic segments were calculated from normalized WGS read depth using JaRMS v0.0.13 as previously described^57^. As JaRMS estimates of genomic copy number are distributed around a value of “1” as the normal diploid copy number count, we multiplied the “levels” estimates from the JaRMS program by two to obtain the adjusted copy number state of genomic regions. JaRMS copy number estimates were used to estimate the population differentiation of taurine and indicine cattle on a per-gene basis using the V_st_ metric^13, 14^. A custom script (CalculateVstDifferences.py) was used to automate the calculation of V_st_ and generation of data tables for plotting. Genes that had a V_st_ higher than 0.3, which is equivalent to the top 1% V_st_, and a difference in average copy number between groups greater than three were considered have a significant difference in copy number between taurine and indicine populations.

In addition to using short WGS reads from the 38 individuals of seven breeds to find structural variants, the haplotype-resolved Angus and Brahman genomes were aligned with the high-quality ARS-UCD1.2 cattle reference to assess structural variants. The advantage of aligning to ARS-UCD1.2 was to standardize the structural variants specific to each haplotype on a common coordinate system. Contigs obtained by breaking final scaffolds at gap positions from UOA_Angus_1 and UOA_Brahman_1 were aligned using nucmer v4^58^ to the ARS-UCD1.2 assembly to identify the larger structural differences (50 bp to 10,000 bp) using Assemblytics^12^. The nucmer alignment parameters were “--maxmatch -t 4 -l 100 -c 500”, which was followed by delta-filter with the option “-g”. Assemblytics parameters followed the default settings, which were “Unique sequence length required: 10000, Maximum variant size: 10000, Minimum variant size: 50”. The overlap of structural variants with Ensembl annotation of Hereford cattle ARS-UCD1.2 release 96 were identified with GenomicFeatures and systemPipeR R packages.

### Identification, copy number and phylogenetic tree of FADS2P1

All chromosomes from Brahman were aligned to the corresponding Angus chromosomes using the dot plot tool Gepard v1.4^59^. Genomic regions that differed between the two subspecies were isolated for further scrutiny. Of all the regions analysed, one particular locus on Brahman chromosome 15 at position ∼4 Mb covering ∼200 kb diverged from the corresponding Angus chromosome. Further analysis revealed an extra copy of fatty acid desaturase 2-like protein (*FADS2P1*) in the Brahman genome. BLASTP^60^ analysis identified two copies of *FADS2P1* in Angus, Hereford, water buffalo and goat, whereas only Brahman had three copies of this gene. A maximum likelihood tree with 1000 bootstraps was constructed for *FADS2P1* homologs using RAxML v8^61^ with substitution model “PROTGAMMAAUTO”. The conservation of synteny around the FADS2P1 locus was investigated by alignments of Angus to Brahman and Angus to Hereford using nucmer v4^58^ and displayed with Ribbon^62^.

### Positive selection analysis on *FADS2P1*

The observation of an indicus specific *FADS2P1* residing in a divergent region prompted further investigation into the possibility that the gene is under positive selection. Homologs of *FADS2P1* in Brahman, Angus and water buffalo were subjected to CODEML analysis as implemented in PAML v4.8^63^. Selective pressure acting on a gene can be estimated by the rate ratio (ω) of non-synonymous (amino acid changes) to synonymous (silent changes) substitutions. Detection of ω > 1 is a sign of positive selection and the site models, namely M7 and M8 in PAML, which allow ω to vary among sites were used to detect positive selection. Protein sequences of *FADS2P1* homologs were aligned using Muscle^64^ and the corresponding nucleotides were mapped back onto the amino acid alignment using PAL2NAL^65^ with gap removal. The tree topology used to run CODEML was a maximum likelihood gene tree calculated from RAxML^61^. Model M8 was compared with M7 using the likelihood ratio test (LRT) to evaluate if the model with positive selection was favoured. More detail on similar positive selection methodology can be found in our study on mammalian GSTs^66^.

### Identification of selective sweep regions

To uncover genetic variants involved in indicine adaptative selection, we designed a strategy to identify selective sweeps using the haplotype-resolved Brahman genome. This method is analogous to the Cross Population Extended Haplotype Homozygosity (XP-EHH)^67^ in that genomic regions are searched for selected alleles that are approaching fixation in the Brahman population but remains polymorphic in six other taurine breeds populations. Only SNPs from the 38 individuals representing seven breeds from the USMARC Beef Diversity Panel version 2.9 (MBCDPv2.9) were considered in the selective sweep analysis. Details are given in Supplementary Note 6. Candidate selective sweep genes in the Brahman genome were analyzed for potential genes overrepresentation in biological pathway using PANTHER 14.1^68^. As there are no *Bos taurus indicus* specific biological pathways annotated, the candidate Brahman genes were mapped to the corresponding Hereford ARS-UCD1.2 Ensembl annotation release96 prior to running PANTHER. The selective sweep intervals were also searched against cattle quantitative trait loci (QTL) from Animal QTL database^69^.

## Statistical analysis

R/Bioconductor was used for all statistical analyses. Significance of positively selected sites found in *FADS2P1* were evaluated using the likelihood ratio test (LRT), with the test statistic *t_LR_* = 2[*l*(*Model* 8)−*l*(*Model* 7)].

## Code availability

Custom scripts can be found at GitHub repository at the following URL: (https://github.com/lloydlow/BrahmanAngusAssemblyScripts)

## Data availability

The PacBio reads, Hi-C reads, RNA-Seq, Iso-Seq and Illumina paired-end reads are available in the SRA under BioProject PRJNA432857. The 38 individuals from seven breeds used for variant calls were downloaded from the BioProject PRJNA324822. The assemblies ARS-UCD1.2 (GCF_002263795.1), Bos_taurus_UMD_3.1.1 (GCF_000003055.6), ARS1 (GCF_001704415.1) and UOA_WB_1 (GCF_003121395.1) were downloaded from the NCBI. Intermediary assembly FASTA files and other miscellaneous information are available from the corresponding authors upon request. Annotation files of UOA_Angus_1 and UOA_Brahman_1 are available through Ensembl.

## URLs

ArrowGrid, https://github.com/skoren/ArrowGrid; seqtk, https://github.com/lh3/seqtk; RepeatMasker, http://www.repeatmasker.org;

## Accessions

**Primary accessions**

BioProject

PRJNA432857

GenBank assembly accession for UOA_Angus_1

GCA_003369685.2

GenBank assembly accession for UOA_Brahman_1

GCA_003369695.2

## Acknowledgements

This work was supported with supercomputing resources provided by the Phoenix HPC service at the University of Adelaide. The work was part funded by the JS Davies bequest to the University of Adelaide. We thank Bob Lee, Kristen Kuhn, Kelsey McClure, and William Thompson for technical assistance. The work was supported in part by funds from USDA-ARS Project Number 3040-31320-012-00D. The use of trade names or commercial products in this manuscript is solely for the purpose of providing specific information and does not imply recommendation or endorsement by the U.S. Department of Agriculture. USDA is an equal opportunity employer and provider. SK, AR, and AMP were supported by the Intramural Research Program of the National Human Genome Research Institute, National Institutes of Health. This work utilized the computational resources of the NIH HPC Biowulf cluster (https://hpc.nih.gov).

## Author contributions

JLW, TPLS, AMP and SH conceived and managed the project; TPLS generated long- and short-read genomic data, as well as RNA-Seq and Iso-Seq data; WYL analysed all results; RT and CL validated sex chromosomes and provided guidance on gene expression work; SK, AR, DMB, BDR, ZNK, SBK, JG and MPH were involved in genome assembly and scaffolding; ARH, JL, and AP provided optical map data; DMB performed CNV analysis; ET provided Iso-Seq FLNC and IsoPhase analysis; FTN, FJM, and KB annotated the genomes; WYL and JLW drafted the manuscript and all authors read, edited, and approved the final manuscript.

## Materials and Correspondence

Correspondence to John L. Williams.

## Supplementary Information

See attached supplementary file.

