## Supplementary Information for "Haplotype-Resolved Cattle Genomes Provide Insights Into Structural Variation and Adaptation"

### Supplementary Figures

Supplementary Figure 1: **Distribution of unassigned PacBio WGS read length.**

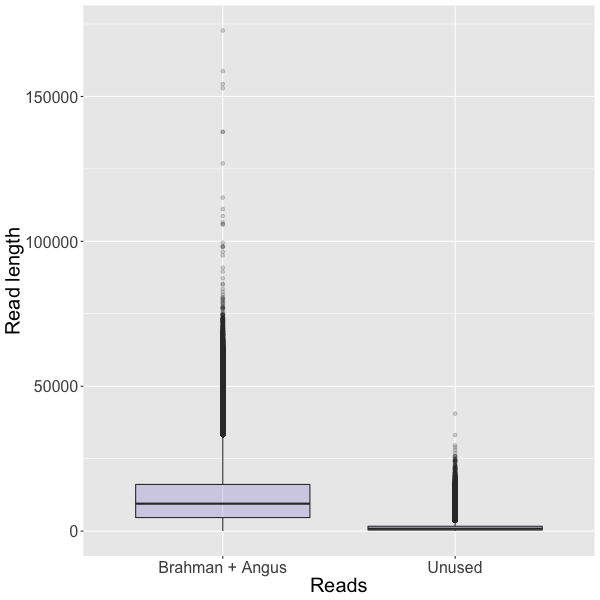

Supplementary Figure 2: **Comparison of chromosome sizes between Angus, Brahman and Hereford assemblies.**

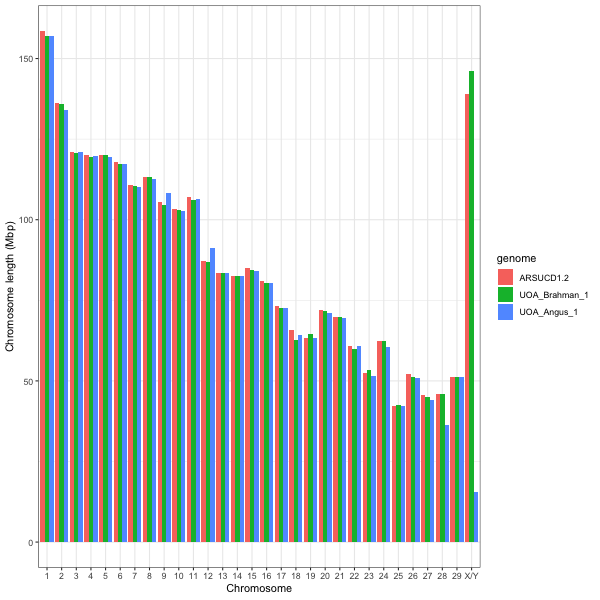

Supplementary Figure 3: **Distribution of un-gapped contig lengths in the three cattle breeds (UOA_Angus_1, UOA_Brahman_1 and ARS-UCD1.2) and water buffalo (UOA_WB_1).**

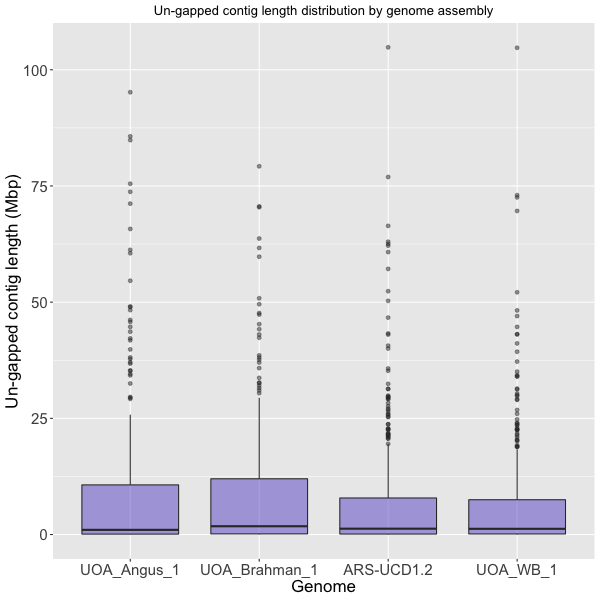

Supplementary Figure 4: **The count in (log_10_ scale) of LINE/L1, LINE/RTE-BovB and Satellite/centromeric repeats in cattle genome assemblies.**

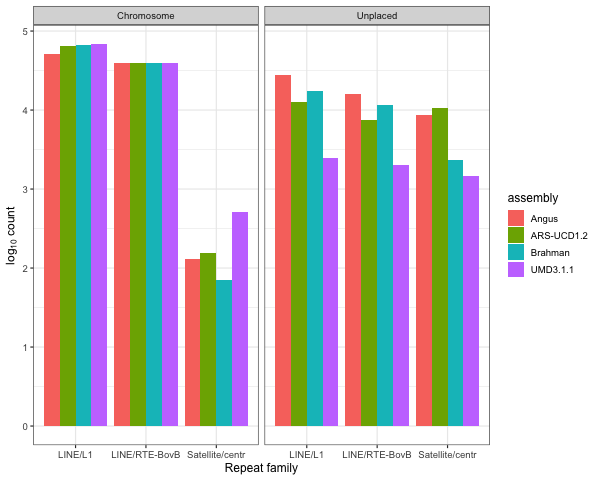

Supplementary Figure 5: **Coverage plot of *FADS2P1* in individuals of different cattle breeds.** The dashed blue line indicates the expected haploid coverage. As *FADS2P1* is member of a gene family, short reads that belong to other gene family members could potentially have mis-mapped to this region, which explains the non-zero coverage in taurine breeds at certain positions across the gene.

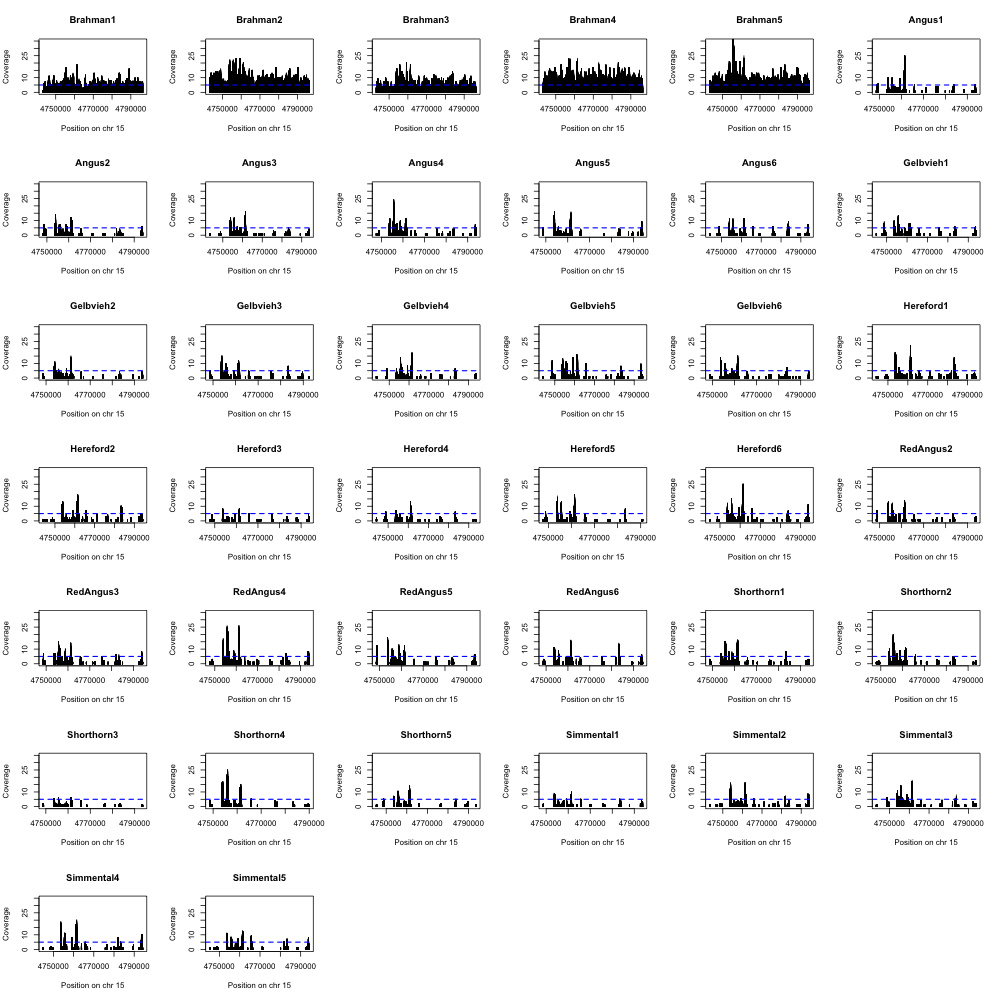

Supplementary Figure 6: **Distribution and breed** **specificity of Brahman and Angus structural variants.** a) Count of structural variants (SVs) categorized as deletion, insertion, repeat contraction, repeat expansion, tandem contraction, and tandem expansion by Assemblytics. The Hereford ARS-UCD1.2 was used as the common reference to call SVs in both Brahman and Angus contigs. b) Venn diagrams showing overlap of six classes of SVs between Brahman and Angus.

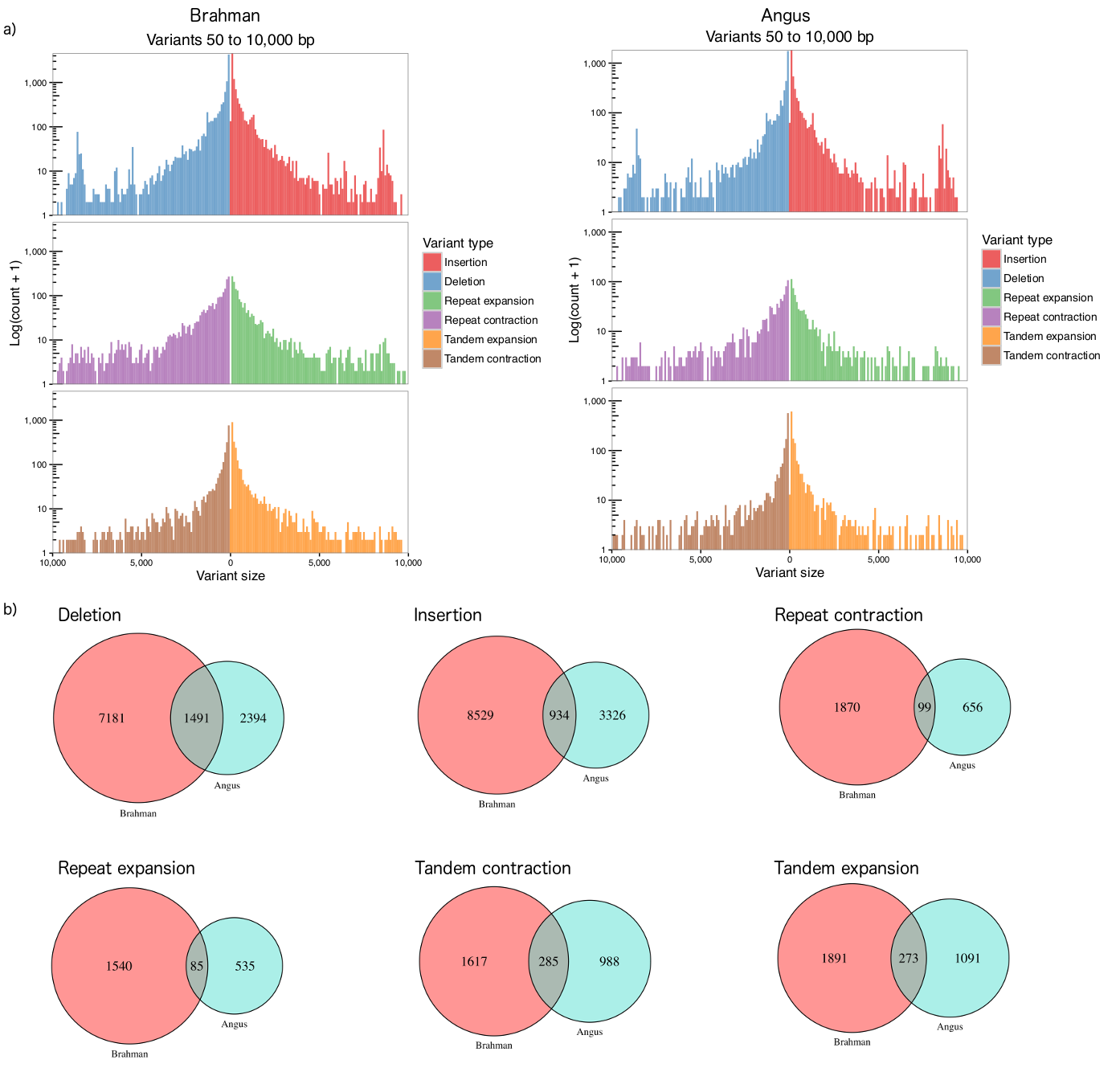

Supplementary Figure 7: **Analysis** **of** **copy number variations using different reference assemblies.** Population differentiation for copy number variations (CNV) as estimated by V_ST_ along each chromosome for the taurine and indicine comparison using a) UOA_Angus_1 and b) ARS-UCD1.2 as the reference genome.

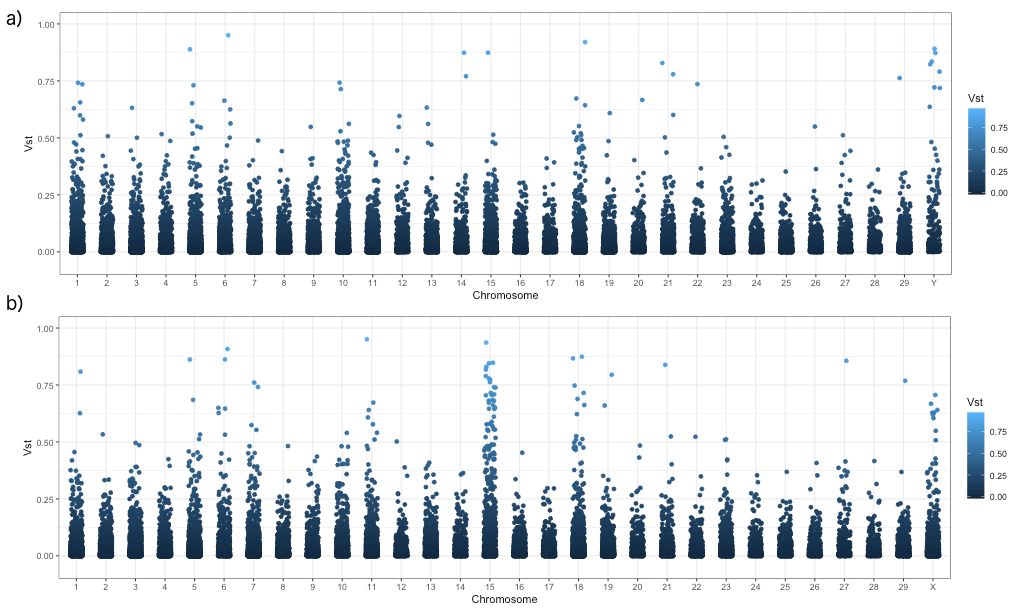

Supplementary Figure 8: **Full-length Iso-Seq transcripts (bottom) and RNA-Seq coverage for *ARIH2* in the brain tissue (top).**

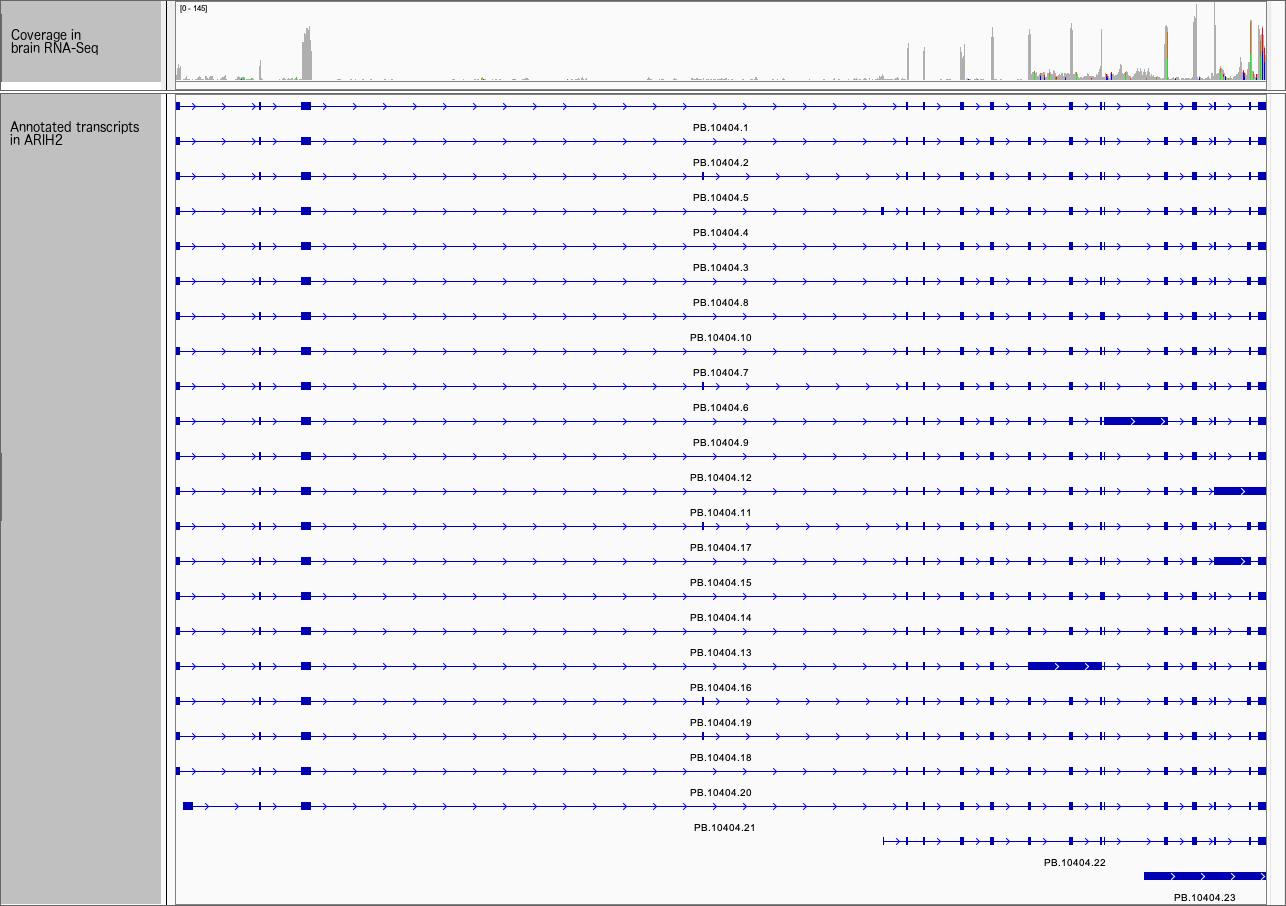

Supplementary Figure 9: **Normalized tissue-specific transcript counts for genes with allelic imbalance and higher expression of the Brahman allele in brain.** a) Calmodulin. b) Pregnancy-associated glycoprotein 1 (*PAG1*).

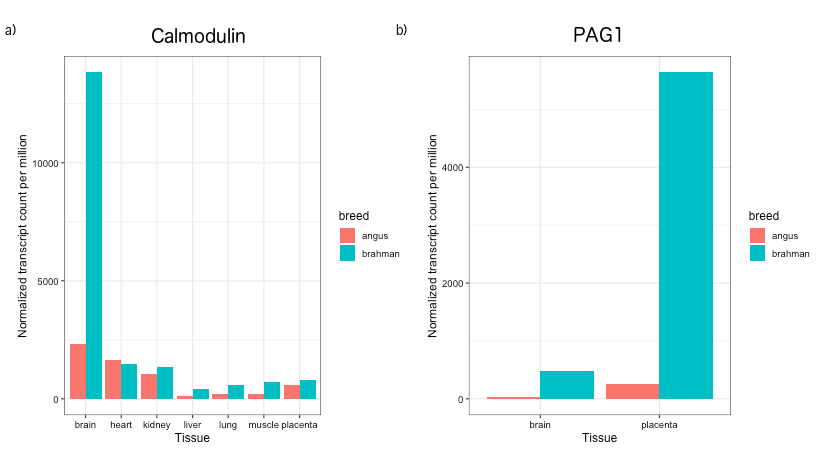

### Supplementary Tables

Supplementary Table 1: **Annotation features in Brahman, Angus and Hereford assemblies.** Comparison of UOA_Brahman_1, UOA_Angus_1, and ARS-UCD1.2 features annotated by the EMBL-EBI pipeline release 96. The columns with X and Y next to each breed show the annotated features in sex chromosomes of the corresponding assemblies.

| Feature | Brahman | Brahman X | Angus | Angus Y | Hereford | Hereford X |
| --- | --- | --- | --- | --- | --- | --- |
| gene | 29910 | 1363 | 28950 | 192 | 27570 | 1132 |
| lncRNA | 3377 | 153 | 3294 | 25 | 1480 | 49 |
| miRNA | 847 | 71 | 868 | 1 | 951 | 73 |
| misc RNA | 386 | 25 | 359 | 0 | 372 | 16 |
| processed pseudogene | 102 | 13 | 97 | 2 | 99 | 7 |
| protein coding | 22118 | 948 | 21419 | 153 | 21848 | 845 |
| pseudogene | 496 | 38 | 454 | 3 | 393 | 29 |
| rRNA | 424 | 13 | 394 | 2 | 393 | 11 |
| ribozyme | 7 | 0 | 6 | 0 | 8 | 1 |
| sRNA | 3 | 0 | 3 | 0 | 3 | 0 |
| scaRNA | 33 | 1 | 31 | 0 | 33 | 1 |
| snRNA | 1195 | 65 | 1146 | 5 | 1201 | 64 |
| snoRNA | 784 | 36 | 752 | 1 | 770 | 36 |

Supplementary Table 2: **Assembly quality score values.**

| Statistic | Description | Angus | Brahman |
| --- | --- | --- | --- |
| QV | Quality value | 44.63 | 46.38 |
| COMPR_PE | Low CE-statistics computed on PE reads | 211314 | 211783 |
| STRECH_PE | High CE-statistics computed on MP reads | 106768 | 92494 |
| LOW_COV_PE | Low read coverage areas | 61490 | 81218 |
| LOW_NORM_COV_PE | Low paired-read coverage areas | 58223 | 79034 |
| HIGH_COV_PE | High read coverage areas | 4329 | 3920 |
| HIGH_NORM_COV_PE | High paired-read coverage areas | 3803 | 2775 |
| HIGH_SPAN_PE | High number of PE reads with pair mapped in a different scaffold | 1808 | 2464 |
| HIGH_SINGLE_PE | High number of PE reads with unmapped pair | 30 | 96 |
| HIGH_OUTIE_PE | High number of mis-oriented or too distant PE reads | 15 | 10 |

Note: CE, compression/expansion; PE, paired-end

Supplementary Table 3: **BUSCO assessment of the completeness of single-copy orthologs for Angus and Brahman genomes.**

| Description | Angus | Brahman |
| --- | --- | --- |
| Complete BUSCOs | 3813 | 3839 |
| Complete and single-copy BUSCOs | 3764 | 3790 |
| Complete and duplicated BUSCOs | 49 | 49 |
| Fragmented BUSCOs | 130 | 123 |
| Missing BUSCOs | 161 | 142 |
| Total BUSCO groups searched | 4104 | 4104 |
| BUSCO completeness (%) | 92.9 | 93.5 |

Supplementary Table 4: **Site models of CODEML for *FADS2P1* and positively selected sites.**

| Model | Log-likelihood | 2Δ(In L) | P-value^b^ | Positively selected sites^a^ | Tree length | Average dN | Average dS |
| --- | --- | --- | --- | --- | --- | --- | --- |
| M7 | -3149.50 |  |  |  |  |  |  |
| M8 | -3132.38 | 17.12 | $0.00019$ | 237V; 271V; 294H; 305C; 306T; 307V**; 311L; 312F; 315V; 317L; 324A; 327C**; 328R; 329R; 330S; 370P | 0.5777 | 0.0168 | 0.0193 |

^a^Bayes Empirical Bayes (BEB) was used to calculate posterior probabilities and only those with$Prob\left( \omega>1 \right)>0.95$ are shown. ** indicates those with $Prob\left( \omega>1 \right)>0.99$. Amino acid position follows Brahman ENSBIXP00005018486.1, which is the indicus-specific copy of *FADS2P1*.

Supplementary Table 5: **Genes identified in the selective sweep intervals.**

| chr | start | end | Ensembl ID | Name | biotype | Indicine mean proportion alternate allele | Taurine mean proportion alternate allele |
| --- | --- | --- | --- | --- | --- | --- | --- |
| 1 | 3500000 | 3600000 | ENSBIXG00005005115 | SCAF4 | protein coding | 0.048 | 0.401 |
| 1 | 81300000 | 81400000 | ENSBIXG00005003979 | SENP2 | protein coding | 0.093 | 0.405 |
| 1 | 81300000 | 81400000 | ENSBIXG00005023240 | LIPH | protein coding | 0.093 | 0.405 |
| 1 | 81300000 | 81400000 | ENSBIXG00005023174 | RF00001 | rRNA | 0.093 | 0.405 |
| 1 | 107100000 | 107200000 | ENSBIXG00005012541 | not available | protein coding | 0.100 | 0.584 |
| 1 | 136400000 | 136500000 | ENSBIXG00005025046 | ACAD11 | protein coding | 0.073 | 0.482 |
| 1 | 136400000 | 136500000 | ENSBIXG00005024967 | DNAJC13 | protein coding | 0.073 | 0.482 |
| 2 | 24300000 | 24400000 | ENSBIXG00005005621 | HAT1 | protein coding | 0.092 | 0.410 |
| 2 | 24300000 | 24400000 | ENSBIXG00005027305 | not available | protein coding | 0.092 | 0.410 |
| 2 | 24300000 | 24400000 | ENSBIXG00005027277 | SLC25A12 | protein coding | 0.092 | 0.410 |
| 3 | 600000 | 700000 | ENSBIXG00005024597 | GPR161 | protein coding | 0.100 | 0.483 |
| 3 | 600000 | 700000 | ENSBIXG00005024283 | DCAF6 | protein coding | 0.100 | 0.483 |
| 3 | 600000 | 700000 | ENSBIXG00005004103 | RF00201 | snoRNA | 0.100 | 0.483 |
| 3 | 600000 | 700000 | ENSBIXG00005023946 | not available | protein coding | 0.100 | 0.483 |
| 3 | 53800000 | 53900000 | ENSBIXG00005030064 | LRRC8C | protein coding | 0.100 | 0.447 |
| 3 | 53800000 | 53900000 | ENSBIXG00005030089 | LRRC8B | protein coding | 0.100 | 0.447 |
| 4 | 400000 | 500000 | ENSBIXG00005018118 | ESYT2 | protein coding | 0.092 | 0.563 |
| 4 | 400000 | 500000 | ENSBIXG00005018040 | NCAPG2 | protein coding | 0.092 | 0.563 |
| 4 | 1000000 | 1100000 | ENSBIXG00005002902 | not available | protein coding | 0.064 | 0.459 |
| 4 | 52900000 | 53000000 | ENSBIXG00005014470 | not available | protein coding | 0.043 | 0.409 |
| 4 | 72600000 | 72700000 | ENSBIXG00005028158 | CDHR3 | protein coding | 0.020 | 0.548 |
| 4 | 72600000 | 72700000 | ENSBIXG00005004834 | not available | miRNA | 0.020 | 0.548 |
| 4 | 95100000 | 95200000 | ENSBIXG00005018443 | CRPPA | protein coding | 0.083 | 0.474 |
| 5 | 42800000 | 42900000 | ENSBIXG00005004767 | FGD4 | protein coding | 0.070 | 0.537 |
| 5 | 42800000 | 42900000 | ENSBIXG00005027380 | RF00026 | snRNA | 0.070 | 0.537 |
| 5 | 48900000 | 49000000 | ENSBIXG00005023898 | SYN3 | protein coding | 0.091 | 0.591 |
| 5 | 106500000 | 106600000 | ENSBIXG00005000567 | not available | protein coding | 0.100 | 0.518 |
| 6 | 116700000 | 116800000 | ENSBIXG00005005681 | not available | protein coding | 0.070 | 0.409 |
| 6 | 116700000 | 116800000 | ENSBIXG00005008386 | not available | lncRNA | 0.070 | 0.409 |
| 6 | 116700000 | 116800000 | ENSBIXG00005008383 | not available | protein coding | 0.070 | 0.409 |
| 6 | 116700000 | 116800000 | ENSBIXG00005000173 | not available | lncRNA | 0.070 | 0.409 |
| 6 | 116700000 | 116800000 | ENSBIXG00005008365 | FGFRL1 | protein coding | 0.070 | 0.409 |
| 6 | 116700000 | 116800000 | ENSBIXG00005008347 | not available | lncRNA | 0.070 | 0.409 |
| 6 | 116700000 | 116800000 | ENSBIXG00005008330 | IDUA | protein coding | 0.070 | 0.409 |
| 7 | 50000000 | 50100000 | ENSBIXG00005019655 | not available | lncRNA | 0.040 | 0.437 |
| 7 | 50000000 | 50100000 | ENSBIXG00005003205 | IL17B | protein coding | 0.040 | 0.437 |
| 7 | 50000000 | 50100000 | ENSBIXG00005003199 | PCYOX1L | protein coding | 0.040 | 0.437 |
| 7 | 50000000 | 50100000 | ENSBIXG00005019586 | not available | protein coding | 0.040 | 0.437 |
| 7 | 50000000 | 50100000 | ENSBIXG00005003178 | AFAP1L1 | protein coding | 0.040 | 0.437 |
| 7 | 50000000 | 50100000 | ENSBIXG00005019455 | not available | protein coding | 0.040 | 0.437 |
| 7 | 51900000 | 52000000 | ENSBIXG00005003009 | JAKMIP2 | protein coding | 0.100 | 0.405 |
| 7 | 66100000 | 66200000 | ENSBIXG00005000684 | HSPA4 | protein coding | 0.100 | 0.459 |
| 7 | 81100000 | 81200000 | ENSBIXG00005020813 | not available | lncRNA | 0.085 | 0.470 |
| 8 | 7300000 | 7400000 | ENSBIXG00005029063 | DEFB136 | protein coding | 0.014 | 0.414 |
| 8 | 7300000 | 7400000 | ENSBIXG00005029055 | CTSB | protein coding | 0.014 | 0.414 |
| 8 | 7300000 | 7400000 | ENSBIXG00005029005 | FDFT1 | protein coding | 0.014 | 0.414 |
| 8 | 10800000 | 10900000 | ENSBIXG00005027580 | not available | protein coding | 0.100 | 0.431 |
| 8 | 10800000 | 10900000 | ENSBIXG00005027560 | ESCO2 | protein coding | 0.100 | 0.431 |
| 8 | 10800000 | 10900000 | ENSBIXG00005004734 | CCDC25 | protein coding | 0.100 | 0.431 |
| 8 | 52900000 | 53000000 | ENSBIXG00005006324 | VPS13A | protein coding | 0.040 | 0.495 |
| 8 | 57500000 | 57600000 | ENSBIXG00005011007 | not available | protein coding | 0.055 | 0.730 |
| 8 | 85000000 | 85100000 | ENSBIXG00005000168 | not available | pseudogene | 0.053 | 0.433 |
| 9 | 32400000 | 32500000 | ENSBIXG00005010729 | CEP85L | protein coding | 0.086 | 0.447 |
| 9 | 40600000 | 40700000 | ENSBIXG00005008671 | FIG4 | protein coding | 0.044 | 0.468 |
| 9 | 40600000 | 40700000 | ENSBIXG00005008574 | AK9 | protein coding | 0.044 | 0.468 |
| 9 | 85800000 | 85900000 | ENSBIXG00005003851 | SASH1 | protein coding | 0.083 | 0.475 |
| 10 | 31600000 | 31700000 | ENSBIXG00005004621 | not available | protein coding | 0.092 | 0.501 |
| 10 | 31600000 | 31700000 | ENSBIXG00005004584 | CAPN3 | protein coding | 0.092 | 0.501 |
| 10 | 72600000 | 72700000 | ENSBIXG00005007972 | GPHN | protein coding | 0.033 | 0.458 |
| 10 | 102100000 | 102200000 | ENSBIXG00005017943 | TTC7B | protein coding | 0.100 | 0.478 |
| 10 | 102100000 | 102200000 | ENSBIXG00005002910 | RF00614 | snoRNA | 0.100 | 0.478 |
| 11 | 37900000 | 38000000 | ENSBIXG00005017832 | CFAP36 | protein coding | 0.100 | 0.498 |
| 11 | 37900000 | 38000000 | ENSBIXG00005017770 | PPP4R3B | protein coding | 0.100 | 0.498 |
| 11 | 37900000 | 38000000 | ENSBIXG00005017706 | PNPT1 | protein coding | 0.100 | 0.498 |
| 11 | 45700000 | 45800000 | ENSBIXG00005001920 | NCK2 | protein coding | 0.075 | 0.406 |
| 11 | 45700000 | 45800000 | ENSBIXG00005001916 | not available | lncRNA | 0.075 | 0.406 |
| 11 | 45700000 | 45800000 | ENSBIXG00005014366 | not available | lncRNA | 0.075 | 0.406 |
| 11 | 45700000 | 45800000 | ENSBIXG00005014363 | RF00619 | snRNA | 0.075 | 0.406 |
| 11 | 45700000 | 45800000 | ENSBIXG00005006127 | not available | lncRNA | 0.075 | 0.406 |
| 11 | 45800000 | 45900000 | ENSBIXG00005007270 | TTL | protein coding | 0.100 | 0.438 |
| 13 | 59700000 | 59800000 | ENSBIXG00005022377 | PIP4K2A | protein coding | 0.060 | 0.441 |
| 14 | 46600000 | 46700000 | ENSBIXG00005016564 | EXT1 | protein coding | 0.087 | 0.432 |
| 14 | 47000000 | 47100000 | ENSBIXG00005007032 | MED30 | protein coding | 0.022 | 0.461 |
| 14 | 64800000 | 64900000 | ENSBIXG00005011378 | VPS13B | protein coding | 0.092 | 0.413 |
| 14 | 64800000 | 64900000 | ENSBIXG00005011322 | RF00156 | snoRNA | 0.092 | 0.413 |
| 15 | 68600000 | 68700000 | ENSBIXG00005002944 | AMOTL1 | protein coding | 0.085 | 0.499 |
| 16 | 4300000 | 4400000 | ENSBIXG00005010567 | MAPKAPK2 | protein coding | 0.085 | 0.473 |
| 16 | 4300000 | 4400000 | ENSBIXG00005010507 | not available | protein coding | 0.085 | 0.473 |
| 16 | 36100000 | 36200000 | ENSBIXG00005023955 | ATP1B1 | protein coding | 0.082 | 0.495 |
| 16 | 36100000 | 36200000 | ENSBIXG00005023904 | RF00155 | snoRNA | 0.082 | 0.495 |
| 16 | 36100000 | 36200000 | ENSBIXG00005023880 | NME7 | protein coding | 0.082 | 0.495 |
| 16 | 36100000 | 36200000 | ENSBIXG00005004083 | not available | protein coding | 0.082 | 0.495 |
| 16 | 48600000 | 48700000 | ENSBIXG00005016454 | not available | protein coding | 0.073 | 0.543 |
| 16 | 48600000 | 48700000 | ENSBIXG00005002621 | DFFB | protein coding | 0.073 | 0.543 |
| 16 | 48600000 | 48700000 | ENSBIXG00005016419 | CEP104 | protein coding | 0.073 | 0.543 |
| 16 | 48600000 | 48700000 | ENSBIXG00005002608 | not available | miRNA | 0.073 | 0.543 |
| 16 | 48600000 | 48700000 | ENSBIXG00005016351 | LRRC47 | protein coding | 0.073 | 0.543 |
| 16 | 48600000 | 48700000 | ENSBIXG00005016324 | SMIM1 | protein coding | 0.073 | 0.543 |
| 16 | 48600000 | 48700000 | ENSBIXG00005016305 | not available | protein coding | 0.073 | 0.543 |
| 16 | 54600000 | 54700000 | ENSBIXG00005008560 | RC3H1 | protein coding | 0.091 | 0.430 |
| 16 | 54800000 | 54900000 | ENSBIXG00005000199 | RABGAP1L | protein coding | 0.035 | 0.447 |
| 16 | 79400000 | 79500000 | ENSBIXG00005019828 | PPP1R12B | protein coding | 0.050 | 0.423 |
| 16 | 79400000 | 79500000 | ENSBIXG00005019795 | RF00004 | snRNA | 0.050 | 0.423 |
| 19 | 27100000 | 27200000 | ENSBIXG00005010360 | not available | protein coding | 0.092 | 0.403 |
| 19 | 27100000 | 27200000 | ENSBIXG00005010357 | MIS12 | protein coding | 0.092 | 0.403 |
| 19 | 27100000 | 27200000 | ENSBIXG00005010333 | DERL2 | protein coding | 0.092 | 0.403 |
| 19 | 27100000 | 27200000 | ENSBIXG00005010287 | DHX33 | protein coding | 0.092 | 0.403 |
| 19 | 27100000 | 27200000 | ENSBIXG00005010261 | NUP88 | protein coding | 0.092 | 0.403 |
| 19 | 27100000 | 27200000 | ENSBIXG00005010218 | RPAIN | protein coding | 0.092 | 0.403 |
| 19 | 27100000 | 27200000 | ENSBIXG00005001571 | RABEP1 | protein coding | 0.092 | 0.403 |
| 19 | 46800000 | 46900000 | ENSBIXG00005022992 | CRHR1 | protein coding | 0.075 | 0.468 |
| 20 | 18500000 | 18600000 | ENSBIXG00005013524 | not available | protein coding | 0.050 | 0.444 |
| 20 | 18500000 | 18600000 | ENSBIXG00005013520 | RF02160 | misc RNA | 0.050 | 0.444 |
| 20 | 18500000 | 18600000 | ENSBIXG00005013509 | RF02159 | misc RNA | 0.050 | 0.444 |
| 21 | 7800000 | 7900000 | ENSBIXG00005023285 | not available | lncRNA | 0.090 | 0.401 |
| 21 | 7800000 | 7900000 | ENSBIXG00005023277 | not available | lncRNA | 0.090 | 0.401 |
| 21 | 7800000 | 7900000 | ENSBIXG00005023256 | IGF1R | protein coding | 0.090 | 0.401 |
| 21 | 49000000 | 49100000 | ENSBIXG00005025684 | SEC23A | protein coding | 0.056 | 0.571 |
| 21 | 49000000 | 49100000 | ENSBIXG00005004365 | GEMIN2 | protein coding | 0.056 | 0.571 |
| 21 | 49000000 | 49100000 | ENSBIXG00005025542 | not available | protein coding | 0.056 | 0.571 |
| 21 | 49000000 | 49100000 | ENSBIXG00005004347 | TRAPPC6B | protein coding | 0.056 | 0.571 |
| 21 | 49000000 | 49100000 | ENSBIXG00005025501 | PNN | protein coding | 0.056 | 0.571 |
| 23 | 14700000 | 14800000 | ENSBIXG00005010033 | KIF6 | protein coding | 0.094 | 0.488 |
| 23 | 43400000 | 43500000 | ENSBIXG00005028562 | not available | lncRNA | 0.083 | 0.573 |
| 24 | 200000 | 300000 | ENSBIXG00005023903 | RF00001 | rRNA | 0.055 | 0.702 |
| 24 | 500000 | 600000 | ENSBIXG00005006840 | PARD6G | protein coding | 0.064 | 0.544 |
| 24 | 600000 | 700000 | ENSBIXG00005004059 | ADNP2 | protein coding | 0.073 | 0.587 |
| 24 | 600000 | 700000 | ENSBIXG00005023670 | not available | protein coding | 0.073 | 0.587 |
| 24 | 600000 | 700000 | ENSBIXG00005023663 | RBFA | protein coding | 0.073 | 0.587 |
| 24 | 37200000 | 37300000 | ENSBIXG00005013581 | SMCHD1 | protein coding | 0.033 | 0.507 |
| 24 | 37200000 | 37300000 | ENSBIXG00005013554 | EMILIN2 | protein coding | 0.033 | 0.507 |
| 24 | 37200000 | 37300000 | ENSBIXG00005013545 | RF00026 | snRNA | 0.033 | 0.507 |
| 27 | 41300000 | 41400000 | ENSBIXG00005026717 | THRB | protein coding | 0.017 | 0.574 |
| 27 | 41300000 | 41400000 | ENSBIXG00005026549 | NR1D2 | protein coding | 0.017 | 0.574 |
| 28 | 18800000 | 18900000 | ENSBIXG00005020285 | not available | protein coding | 0.027 | 0.415 |
| 28 | 18800000 | 18900000 | ENSBIXG00005020283 | ADO | protein coding | 0.027 | 0.415 |
| 28 | 18800000 | 18900000 | ENSBIXG00005020273 | EGR2 | protein coding | 0.027 | 0.415 |
| 29 | 19200000 | 19300000 | ENSBIXG00005028250 | ETS1 | protein coding | 0.100 | 0.458 |

Supplementary Table 6: **Annotation of SNP and INDEL variants.** Short read data from either Angus or Brahman was mapped to the corresponding reference genomes. After GATK variant calling and filtering of variants, Annovar was used to annotate the variants identified. Note: SNV is single nucleotide variant.

| Description | Angus | Brahman |
| --- | --- | --- |
| number of animals | 6 | 5 |
| nonsynonymous SNV | 53730 | 79170 |
| stop gain | 871 | 1253 |
| stop loss | 220 | 267 |
| synonymous SNV | 47843 | 96675 |
| frameshift deletion | 1350 | 1866 |
| frameshift insertion | 1120 | 1397 |
| nonframeshift deletion | 519 | 845 |
| nonframeshift insertion | 386 | 588 |
| stop gain | 61 | 101 |
| stop loss | 9 | 15 |

Supplementary Table 7: **Breed-specific structural variant (SV) type and over/under-represented gene ontology for biological processes.**

| **Angus-specific insertion SV** |  |  |  |  |
| --- | --- | --- | --- | --- |
| PANTHER GO-Slim Biological Process | Over/Under-represented GO | Fold enrichment | Raw P-value | FDR |
| cellular response to stimulus (GO:0051716) | + | 1.75 | 8.49E-06 | 1.52E-02 |
| **Angus-specific tandem contraction SV** |  |  |  |  |
| PANTHER GO-Slim Biological Process | Over/Under-represented GO | Fold enrichment | Raw P-value | FDR |
| synaptic vesicle endocytosis (GO:0048488) | + | 24.92 | 3.75E-06 | 3.35E-03 |
| synaptic vesicle cycle (GO:0099504) | + | 19.29 | 1.14E-05 | 5.08E-03 |
| organophosphate biosynthetic process (GO:0090407) | + | 15.43 | 1.96E-04 | 3.18E-02 |
| peptide metabolic process (GO:0006518) | + | 13 | 6.40E-05 | 1.91E-02 |
| regulation of cation transmembrane transport (GO:1904062) | + | 12.46 | 7.71E-05 | 1.97E-02 |
| regulation of ion transmembrane transport (GO:0034765) | + | 12.46 | 7.71E-05 | 1.72E-02 |
| regulation of ion transport (GO:0043269) | + | 10.47 | 8.03E-06 | 4.78E-03 |
| regulation of transport (GO:0051049) | + | 7.9 | 4.43E-05 | 1.58E-02 |
| membrane invagination (GO:0010324) | + | 6.2 | 1.86E-04 | 3.70E-02 |
| vesicle budding from membrane (GO:0006900) | + | 6.2 | 1.86E-04 | 3.33E-02 |
| regulation of localization (GO:0032879) | + | 6.06 | 3.37E-06 | 6.03E-03 |
| **Brahman-specific insertion SV** |  |  |  |  |
| PANTHER GO-Slim Biological Process | Over/Under-represented GO | Fold enrichment | Raw P-value | FDR |
| release of sequestered calcium ion into cytosol (GO:0051209) | + | 6.93 | 1.87E-04 | 2.78E-02 |
| negative regulation of sequestering of calcium ion (GO:0051283) | + | 6.67 | 2.29E-04 | 2.56E-02 |
| phospholipid translocation (GO:0045332) | + | 5.71 | 8.27E-05 | 2.11E-02 |
| organophosphate biosynthetic process (GO:0090407) | + | 5.59 | 5.72E-04 | 4.09E-02 |
| lipid translocation (GO:0034204) | + | 5.57 | 9.78E-05 | 2.19E-02 |
| sequestering of calcium ion (GO:0051208) | + | 5.57 | 9.78E-05 | 1.94E-02 |
| positive regulation of cell migration (GO:0030335) | + | 5.5 | 2.56E-04 | 2.54E-02 |
| regulation of ion transport (GO:0043269) | + | 3.71 | 2.28E-04 | 2.71E-02 |
| lipid transport (GO:0006869) | + | 3.32 | 3.42E-04 | 2.91E-02 |
| negative regulation of cellular process (GO:0048523) | + | 3.27 | 2.35E-04 | 2.47E-02 |
| microtubule-based movement (GO:0007018) | + | 3.19 | 7.74E-04 | 4.77E-02 |
| phosphate-containing compound metabolic process (GO:0006796) | + | 3.12 | 5.74E-04 | 3.94E-02 |
| regulation of transport (GO:0051049) | + | 3.04 | 7.32E-04 | 4.85E-02 |
| lipid localization (GO:0010876) | + | 3.04 | 7.32E-04 | 4.67E-02 |
| regulation of membrane potential (GO:0042391) | + | 2.96 | 1.56E-04 | 2.54E-02 |
| organophosphate metabolic process (GO:0019637) | + | 2.88 | 2.11E-04 | 2.90E-02 |
| regulation of localization (GO:0032879) | + | 2.74 | 2.50E-05 | 7.44E-03 |
| membrane fusion (GO:0061025) | + | 2.7 | 4.27E-04 | 3.18E-02 |
| microtubule-based process (GO:0007017) | + | 2.61 | 1.04E-04 | 1.86E-02 |
| macromolecule localization (GO:0033036) | + | 2.53 | 4.23E-04 | 3.29E-02 |
| signal transduction (GO:0007165) | + | 1.65 | 2.29E-07 | 1.36E-04 |
| intracellular signal transduction (GO:0035556) | + | 1.64 | 3.88E-04 | 3.15E-02 |
| cellular response to stimulus (GO:0051716) | + | 1.57 | 6.15E-07 | 2.20E-04 |
| localization (GO:0051179) | + | 1.41 | 2.21E-04 | 2.83E-02 |
| cellular process (GO:0009987) | + | 1.27 | 4.24E-07 | 1.90E-04 |
| gene expression (GO:0010467) | - | 0.62 | 3.32E-04 | 2.97E-02 |
| sensory perception (GO:0007600) | - | 0.35 | 2.75E-04 | 2.59E-02 |
| sensory perception of chemical stimulus (GO:0007606) | - | 0.05 | 4.59E-08 | 4.10E-05 |
| detection of chemical stimulus involved in sensory perception (GO:0050907) | - | < 0.01 | 1.12E-08 | 2.00E-05 |
| **Brahman-specific deletion SV** |  |  |  |  |
| PANTHER GO-Slim Biological Process | Over/Under-represented GO | Fold enrichment | Raw P-value | FDR |
| organophosphate biosynthetic process (GO:0090407) | + | 5.9 | 4.21E-04 | 4.18E-02 |
| small molecule biosynthetic process (GO:0044283) | + | 5.5 | 2.49E-04 | 3.43E-02 |
| sequestering of calcium ion (GO:0051208) | + | 5.22 | 3.38E-04 | 4.32E-02 |
| calcium-mediated signaling (GO:0019722) | + | 3.82 | 1.75E-04 | 2.84E-02 |
| cellular calcium ion homeostasis (GO:0006874) | + | 3.22 | 8.44E-06 | 5.03E-03 |
| calcium ion homeostasis (GO:0055074) | + | 3.21 | 9.16E-06 | 4.09E-03 |
| negative regulation of cellular process (GO:0048523) | + | 3.2 | 4.55E-04 | 4.28E-02 |
| divalent inorganic cation homeostasis (GO:0072507) | + | 2.98 | 1.64E-05 | 5.86E-03 |
| ion homeostasis (GO:0050801) | + | 2.43 | 5.87E-05 | 1.31E-02 |
| inorganic ion homeostasis (GO:0098771) | + | 2.37 | 1.10E-04 | 1.97E-02 |
| chemical homeostasis (GO:0048878) | + | 2.35 | 3.85E-05 | 9.82E-03 |
| homeostatic process (GO:0042592) | + | 2.05 | 3.56E-04 | 4.24E-02 |
| regulation of biological quality (GO:0065008) | + | 1.75 | 3.83E-04 | 4.02E-02 |
| intracellular signal transduction (GO:0035556) | + | 1.71 | 1.90E-04 | 2.83E-02 |
| signal transduction (GO:0007165) | + | 1.53 | 2.44E-05 | 7.27E-03 |
| cellular response to stimulus (GO:0051716) | + | 1.4 | 3.62E-04 | 4.05E-02 |
| cellular process (GO:0009987) | + | 1.22 | 7.56E-05 | 1.50E-02 |
| sensory perception of chemical stimulus (GO:0007606) | - | 0.1 | 1.30E-06 | 1.17E-03 |
| detection of chemical stimulus involved in sensory perception (GO:0050907) | - | < 0.01 | 2.33E-08 | 4.17E-05 |

### Supplementary Notes

Supplementary Note 1: **Comparison of different Hi-C scaffolding programs**

Three different scaffolders, 3D-DNA^1^, Proximo (Phase Genomics) and SALSA2^2^ were evaluated for building scaffolds using the following parameters.

For 3D-DNA, raw reads were aligned with juicer git commit d940e9e012a75822ff3f5a9ed7b3ecf08999df01 with the options -z `pwd`/reference/asm.fasta -y `pwd`/reference/asm_MboI.txt -q phillippy.q -l phillippy.q -D software/juicer/ -d `pwd` -p `pwd`/reference/chr.sizes. Scaffolding with 3D de novo assembly: version 170123 used the command -m haploid -t 15000 -s 2 -c 30 asm.fasta merged_nodups.txt for both haplotypes.

Phase Genomics' Proximo Hi-C genome scaffolding platform (git commit 145c01be162be85c060c567d576bb4786496c032) was used to create chromosome-scale scaffolds from the contig assembly as described in Bickhart et al^3^. As in the LACHESIS method^4^, this process computes a contact frequency matrix from the aligned Hi-C read pairs, normalized by the number of Sau3AI restriction sites (GATC) on each contig, and constructs scaffolds in such a way as to optimize expected contact frequency and other statistical patterns in Hi-C data. Approximately 40,000 separate Proximo runs were performed to optimize the number of scaffolds and scaffold construction in order to make the scaffolds as concordant with the observed Hi-C data as possible.

Details for the SALSA2 run is given in the Methods section of the manuscript.

We evaluated the performance of each scaffolder using a heuristic scoring method involving mapping genetic markers to each set of scaffolds. Each method was scored based on the inverse of the number of scaffolds required to cover the base chromosome (N), the difference in cumulative length of these scaffolds compared to the analog chromosome in ARS-UCD1.2 in megabases (L), and the number of contig order errors within each scaffold. The score function can be summarized by this equation:

$$Score= \frac{1}{(N+i+b+L)}$$

where “i” represents the number of contig inversions and “b” represents the number of continuity breaks detected in the genetic map based on their positions in ARS-UCD1.2. Scoring was performed on a per-chromosome basis with replacement of scaffolds that had previously mapped to other chromosomes. Using this method, SALSA2 gave most chromosomes with the highest scores and hence was the chosen to use for scaffolding.

Supplementary Note 2: **Comparison of optical map based scaffolding approaches**

*De novo* optical map assembly and haplotype resolution

Approximately 450 Gb of sequence, representing ~167x coverage, with molecule length >150 kb was aligned to the Brahman and Angus haplotig assemblies respectively to produce haplotype resolved scaffolds of each breed. Alignment of each molecule to the haplotigs of each breed was given a confidence score, which is the log of the alignment p-value. If the confidence score was 2 points higher for one breed haplotigs than the other, then the molecule was binned with the breed with the higher score. Molecule with alignments having almost equal score, defined as within 2 confidence score points of each other, were considered as homozygous and were randomly binned to one of the two breeds to keep the coverage in homozygous and heterozygous regions uniform. In summary, 135 Gb of the molecules aligned to Angus only, 141 Gb of the molecules aligned to Brahman only, 110 Gb aligned to both Angus and Brahman and were binned evenly. About 64 Gb of molecules aligned to neither the Brahman nor the Angus haplotigs and were also binned randomly to one or the other breed.

For the optical map assembly, a pairwise comparison of all DNA molecules was used to create a layout overlap graph, which was then used to create the initial consensus genome maps. By realigning molecules to the genome maps and using only the best-matched molecules, the label positions on the genome maps were refined and used to validate the minimum assembly tiling path. Next, the software aligned molecules to genome maps and extended the maps based on the molecules aligning beyond the map ends. Overlapping genome maps were then merged. This process was repeated 5 times before a final refinement step was applied to “finish” all genome maps.

To analyse the advantages of haplotype-resolved vs haplotype-unaware optical map construction to guide scaffolding, we generated scaffolds based on the conventional approach that used all Bionano molecules. With about 450 Gbp (molecules >150kbp) of molecules collected from the Brahman-Angus offspring, the *de novo* assembly was 3.37 Gbp with an N50 of 71.11 Mbp. This approach was not biased a priori by assigning parental haplotype and instead resolved haplotype in a *de novo* manner. The advantages of this approach were an increased sequence coverage as molecules were not split to each haplotype and there was no reliance on molecule alignment to haplotigs to bin them to each breed. However, as parental alleles were not separated prior to assembly, switching between parental alleles in the scaffolds is possible. Furthermore, as the scaffolding algorithm was unaware that the Angus contigs have no X sequences, the final scaffold length for the Angus assembly was longer than expected as the genome map used to guide scaffolding included the X chromosome. The Angus sequence that aligned to the X chromosome genome map likely belonged to the Y chromosome pseudoautosomal region, which is known to have high sequence identity with the X chromosome.

Supplementary Note 2 Table 1: **Input dataset used to perform optical map-based assemblies using the haplotype-resolved versus the conventional haplotype unaware approach.**

| Description | Angus × Brahman Offspring | Angus-Selected | Brahman-Selected |
| --- | --- | --- | --- |
| Data collected (molecules > 150 kbp) | 480 Gbp | 222 Gbp | 228 Gbp |
| Effective coverage of reference | 126x | 65x | 65x |
| Assembly size | 3.37 Gbp | 2.79 Gbp | 2.87 Gbp |
| Genome map N50 | 71.11 Mbp | 33.97 Mbp | 28.62 Mbp |

Haplotype-resolved scaffold assembly

Haplotype-resolved scaffold assembly was performed using the contigs of Brahman and Angus separately, and the Bionano genome map was assembled using standard parameters in Bionano Access (Bionano Solve 3.2.1). For both the Brahman-selected and Angus-selected assemblies, about 98% of the sequences were incorporated into the final hybrid assemblies with N50s of about 34 Mbp. The Angus-selected scaffold has a length of 2.53 Gbp while the Brahman-selected scaffold has a length of 2.64 Gbp. The N50s of the Angus-selected and Brahman-selected assemblies are limited by the breakage of the Brahman and Angus sequence assemblies in potentially random regions of the genome, which creates a bias in the molecule alignment step during molecule selection. During the scaffolding process, 29 and 36 discrepancies were identified in the Angus and Brahman scaffolds, respectively. These were most likely sequence chimeras, and breaks were introduced in the sequence contigs.

Supplementary Note 2 Table 2: **Angus-selected Bionano assembly with Angus contigs.**

| Statistic | Original Bionano | Original sequence | Sequence used in scaffold | Scaffold | Scaffold + leftover unscaffolded sequence |
| --- | --- | --- | --- | --- | --- |
| Number of contigs | 597 | 1747 | 397 | 217 | 1595 |
| N50 (Mbp) | 33.97 | 29.44 | 32.50 | 35.49 | 35.24 |
| Total length (Mbp) | 2790.07 | 2573.81 | 2512.48 (97.6%) | 2526.00 | 2587.27 |

Supplementary Note 2 Table 3: **Brahman-selected Bionano assembly with Brahman contigs.**

| Statistic | Original Bionano | Original sequence | Sequence used in scaffold | Scaffold | Scaffold + leftover unscaffolded sequence |
| --- | --- | --- | --- | --- | --- |
| Number of contigs | 493 | 1585 | 421 | 154 | 1353 |
| N50 (Mbp) | 28.62 | 23.45 | 21.99 | 32.70 | 31.74 |
| Total length (Mbp) | 2867.64 | 2678.77 | 2632.15 (98.3%) | 2644.13 | 2690.21 |

Conventional optical map scaffold assembly

Scaffolding between the Brahman-Angus offspring Bionano map with the Brahman sequence and the Angus sequence were also performed respectively (Bionano Solve 3.2.2). The assemblies have lengths of about 2.65 Gbp with N50 of 84 Mbp. Potential sequence errors (133 in the Angus and 65 in the Brahman) were detected and corrected while running the Bionano scaffolding pipeline (Bionano Solve 3.2.2).

Supplementary Note 2 Table 4: **Offspring scaffolding with Angus contigs.**

| Statistic | Original Bionano | Original sequence | Sequence used in scaffold | Scaffold | Scaffold + leftover unscaffolded sequence |
| --- | --- | --- | --- | --- | --- |
| Number of contigs | 1026 | 1747 | 496 | 111 | 1414 |
| N50 (Mbp) | 71.11 | 29.44 | 29.44 | 84.07 | 84.07 |
| Total length (Mbp) | 3370.21 | 2573.81 | 2518.93 (97.9%) | 2643.34 | 2693.03 |

Supplementary Note 2 Table 5: **Offspring scaffolding with Brahman contigs.**

| Statistic | Original Bionano | Original sequence | Sequence used in scaffold | Scaffold | Scaffold + leftover unscaffolded sequence |
| --- | --- | --- | --- | --- | --- |
| Number of contigs | 1026 | 1585 | 433 | 87 | 1282 |
| N50 (Mbp) | 71.11 | 23.45 | 21.82 | 84.31 | 84.31 |
| Total length (Mbp) | 3370.21 | 2678.77 | 2632.43 (98.3%) | 2660.03 | 2705.93 |

Supplementary Note 3: **Genome annotation of UOA_Brahman_1 using the NCBI annotation pipeline**

The NCBI Eukaryotic Genome Annotation Pipeline was used to annotate genes, transcripts, proteins and other genomic features on the Bos indicus haplotype (GCF_003369695.1). The methodology for producing NCBI Bos indicus x Bos taurus Annotation Release 100 (AR 100) was as described for the UMD_CASPUR_WB_2.0 assembly^5^. The evidence aligned to the genome and used for gene prediction was made up of transcript data from the same individual that provided the sample for the genomic sequence: 56,550 PacBio Iso-Seq reads from brain, liver, kidney, placenta, skeletal muscle, lung, and heart, and 1.7 billion RNA-Seq reads from brain, liver, lung, placenta and skeletal muscle. The other evidence used were: 8 billion RNA-Seq reads from 15 *Bos indicus* tissue samples, transcripts and proteins from *Bos taurus* (14,281 known RefSeq proteins, 19, 584 GenBank proteins, 1,583,270 ESTs), and 52,350 human known RefSeq proteins.

The resulting annotation consists of 20,846 protein-coding genes, 16,398 of which have an ortholog to human. Only 135 protein-coding genes are missing a start or a stop codon and are marked as partial. Where necessary, the annotation pipeline introduced differences between the predicted models and the genomic sequence to compensate for frameshift-causing genomic insertions or deletions that are not supported by protein alignments. These “corrected” proteins are prefixed with ‘LOW QUALITY PROTEIN’ and should be considered as lower confidence. As an additional testament to the quality of the assembly, only 677 protein coding genes required such a “correction”.

Supplementary Note 3 Table 1: **Comparisons of the number of corrected coding sequences in selected mammalian species.**

| Scientific name | Assembly name | Sequencing technology | Number of corrected CDS |
| --- | --- | --- | --- |
| *Bos taurus* | ARS-UCD1.2 | PacBio; Illumina NextSeq 500; Illumina HiSeq; Illumina GAII | 622 |
| *Bos indicus x Bos taurus* | UOA_Brahman_1 | PacBio Sequel; PacBio RSII; Illumina NextSeq | 677 |
| *Capra hircus* | ARS1 | PacBio | 946 |
| *Ovis aries musimon* | Oori1 | Not listed | 979 |
| *Bos indicus* | Bos_indicus_1.0 | SOLiD | 1383 |
| *Bison Bison* | Bison_UMD1.0 | 454; Illumina HiSeq | 1,448 |
| *Ovis aries* | Oar_rambouillet_v1.0 | HiSeq X Ten; PacBio RS II | 1,646 |
| *Bubalus bubalis* | UOA_WB_1 | PacBio | 1943 |
| *Bos mutus* | BosGru_v2.0 | Illumina HiSeq; Illumina GA | 1954 |
| *Ovis aries* | Oar_v4.0 | Illumina GAII; 454; PacBio RSII | 4524 |

Supplementary Note 4: **Genome annotation of UOA_Angus_1 and UOA_Brahman_1 using the Ensembl annotation pipeline**

Four major classes of evidence were used to create a set of candidate transcripts: pooled high-quality Iso-Seq transcripts from seven tissues (brain, liver, lung, skeletal muscle, placenta, kidney, and heart) of the sequenced F1 hybrid fetus, short read RNA-seq data from five tissues (brain, liver, lung, skeletal muscle, and placenta) of the F1 hybrid fetus, along with publicly available *Bos taurus taurus* and *Bos taurus indicus* data, human transcripts mapped from the GENCODE^6^ gene set using pairwise whole genome alignment and finally vertebrate proteins with experimental evidence from UniProt^7^ (see below).

Supplementary Note 4 Table 1: **Initial transcript models for each major input data type for Ensembl annotation.** The unfiltered transcript models came from the initial alignments of each datatype, and later in the annotation process low quality, fragmented and redundant models were removed to produce the finalized gene/transcript models.

| Data type | Maternal initial models | Paternal  initial models |
| --- | --- | --- |
| Iso-Seq (7 pooled tissues) | 165822 | 161968 |
| Short read RNA-seq (32 tissues + 1 merged) | 1167059 | 1150303 |
| Human GENCODE mapping | 53178 | 52242 |
| UniProt known vertebrate proteins | 542891 | 533051 |

Data from each locus was assessed to remove low quality models and then collapsed into a final gene model with an associated non-redundant transcript set. During the collapsing process, priority was given to transcript models based on the transcriptomic (Iso-Seq and RNA-Seq) data over models derived from homology. For protein coding genes, we also assessed the coverage of the open reading frame (ORF) in relation to known vertebrate proteins. The most complete model was chosen at each locus. In cases where the transcriptomic data appeared fragmented in comparison to the homology data, the homology data were included for completeness. Similarly, for regions where there were no transcriptomic data, we included models based on homology if there was a sufficiently good alignment.

Each gene was then classified as protein coding, long non-coding or pseudogene based on an analysis of the alignment information present at each locus. Genes with transcripts matching known proteins, which did not display multiple structural abnormalities, were classified as protein coding. If a gene matched a known protein but had several problems with the underlying structure (i.e. non-canonical splice sites, abnormally short introns, high level of repeat coverage, no evidence of expression), we classified it as a pseudogene. Single exon genes were assessed for evidence of retrotransposition based on the presence of a multi-exon gene with a highly similar ORF elsewhere in the genome. Such single exon genes were classed as processed pseudogenes. If a gene fell into none of the previous categories, did not overlap a protein coding gene and had been constructed from transcriptomic data, it was considered as a potential lncRNA. The lncRNA set was filtered to remove transcripts that did not have at least two valid splice sites or cover 1000 bp.

In addition to the above, a small non-coding RNA annotation was produced. The miRNA genes were identified by running a BLAST^8^ of miRbase^9^ against the genome and then passing the results into RNAfold^10^. Results were post filtered to remove poor quality alignments or alignments that were covered by repeats. For other small non-coding gene types, Rfam^11^ was used to scan against the genome and the results were passed into Infernal^12^. Both the maternal and paternal gene sets are available as part of Ensembl release 97. More information on the annotation is available at <http://asia.ensembl.org/info/genome/genebuild/2019_06_hybrid_cattle_gene_annotation.pdf>

Supplementary Note 5: **Further assembly evaluation**

The completeness of the genome from contig to chromosome-level assembly was assessed using the Benchmarking Universal Single-Copy Orthologs (BUSCO) v2.0.1^13^. The mammalia_odb9 lineage-specific profile that contains 4,104 BUSCO gene groups was tested against the Brahman and Angus genome assemblies using the option “-m geno”. In addition to testing for completeness in gene space using BUSCO, the assembly base quality values (QV) and other assembly metrics including compression/expansion (CE) errors were calculated using BWA MEM^14^, FRCBam^15^ and Freebayes^16^ as previously described^17^.

For consistency in the evaluation of some recently assembled mammalian genomes, we assessed the error rates using the same method described for the water buffalo^18^ and goat^3^ genomes. Briefly, short-insert Illumina WGS reads from the parents of the sequenced hybrid F1 animal were aligned to the corresponding haplotype-resolved assemblies using BWA MEM^14^. These short reads were not used in the assembly process and hence they served as an independent dataset to evaluate the genomes. We used the reference-free assembly validation software, FRCBam^15^ to generate feature response curves on the Angus and Brahman assemblies in order to identify compression/expansion (CE) errors in the genomes. Putative erroneous bases in each assembly were also identified using FreeBayes^16^. The results are tabulated in Supplementary Table 2. Commands used to generate all assembly quality assessment metrics can be found in the GitHub repository (<https://github.com/lloydlow/BrahmanAngusAssemblyScripts>).

Supplementary Note 6: **The selection sweep method**

If a genomic region has multiple SNPs close to fixation or a high level of homozygosity within Brahman individuals, but the same region contains more segregating polymorphic variants in taurine breeds, it can be considered as a candidate region for a selective sweep. An alternative explanation for such SNP patterns is genetic drift. Individuals chosen for selective sweep detection were sires with minimal pedigree relationships to ensure sampling of diverse alleles suitable for a population genetics study^19^. The methodology for is similar to other recent studies to identifying selective sweep in cattle^20,21^. The present study had the advantage of using a haplotype-resolved Brahman genome to call SNPs rather than relying on the poorer resolution taurine-based reference, UMD3.1.1.

The proportion of alternate alleles in fixed size windows for the 38 animals representing seven breeds was calculated using the Brahman reference genome. To calculate the proportion of alternate alleles, the annotated genotype data was processed in R using custom scripts to ensure good quality data was included. Only genotypes with complete calls for all animals were retained. For Brahman genotypes, a further requirement of at least five mapped reads was imposed to ensure sufficient coverage to detect the presence of alternate alleles. A genotype labelled as 0/0 is homozygous and the same as the reference and hence there is no alternate allele. The maximum number of alternate alleles possible at a locus for an individual is two (e.g. a genotype call of 1/1).

Let

$n=$ Number of individuals

$c=$ Count of alternate allele at a particular position per individual

$m=$ Number of SNPs in the window

Proportion of alternate allele at position $j$ is

$$\frac{\sum_{i=1}^{n} c_{ij}}{2n}$$

where $c_{11}$ refers to the count of alternate allele for the 1^st^ individual at the 1^st^ SNP in the window.

Therefore, for each window, mean proportion of alternate allele (MP) is,

$$MP=\frac{\sum_{j=1}^{m} \left( \frac{\sum_{i=1}^{n} c_{ij}}{2n} \right)}{m}$$

Using this formula, indicine individuals were grouped together to calculate proportion mean of alternate allele, and the same process was repeated for the taurine individuals as a group. The figure below shows the proportion of mean alternate alleles in Brahman and the other six taurine breeds as histograms.

Supplementary Note 6 Figure 1: **Histograms of the mean proportion alternate allele in Brahman and the other six taurine breeds.** The red line to the left in the Brahman plot indicates the bottom 5% percentile (i.e. extremely low polymorphism). The red line to the right in the taurine plot indicates the top 10% percentile with high polymorphic divergence relative to the Brahman.

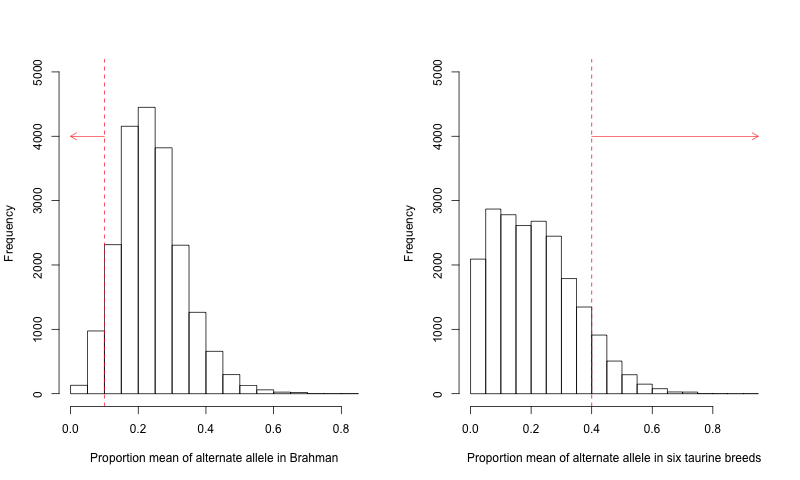

Three fixed size windows (50 kb, 100 kb and 150 kb) were tested to check which would cover sufficient SNPs for the calculation of alternate alleles, and to determine the distance of consecutive SNPs. The table below shows the comparison of different window sizes for the dataset.

Supplementary Note 6 Table 1: **Comparisons of different window sizes for SNP counts and consecutive SNP distances in each window.**

| Window size (kb) | Mean SNP count | Median SNP count | Mean consecutive SNP distance (bp) | Median consecutive SNP distance (bp) | Percentage of windows with at least 10 SNPs |
| --- | --- | --- | --- | --- | --- |
| 50 | 8.381 | 8.000 | 3341 | 3306 | 36% |
| 100 | 16.39 | 16.00 | 5321 | 5028 | 81% |
| 150 | 24.39 | 24.00 | 6122 | 5562 | 92% |

The 50 kb window size was not appropriate for the selective sweeps analysis because many windows contained less than 10 SNPs. When the window size selected was 100 kb, 81% of all windows contained at least 10 SNPs. Additionally, the distance between consecutive SNPs was ~800 bp shorter, on average, when compared to the 150 kb window size. In other words, there was higher density of SNPs when 100 kb windows were chosen. Regardless of whether the 100 kb or the 150 kb window size was chosen, the final gene list in the selection intervals overlapped substantially, although more candidate genes were detected using 100 kb windows. Only windows with at least 10 SNPs were considered for calculation of mean proportion of alternate alleles.

Supplementary Note 7: **Further Iso-Seq analysis**

**SNP calls missed by IsoPhase are either in homopolymer regions or have low coverage**

There are 8,093 (substitution) SNP calls that are missed by IsoPhase but called jointly by RNA-Seq and genomic data. 5589 (69%) of the missed calls are either within or adjacent to a homopolymer (HP) region. IsoPhase defines a HP region as a stretch of 4 consecutive identical bases and will not call a SNP if it is inside a HP or immediately adjacent to a HP region.

Of the remaining 1813 missed calls (22%), have effective base coverage less than 40 in the Iso-Seq data. Effective base coverage is different from full-length Iso-Seq read coverage because after alternative splicing, certain exons, introns, or UTRs may have lower coverage despite meeting the initial 40 fold threshold requirement to run IsoPhase.

Of the missed PacBio calls, 91% are either in or adjacent to HP regions, or have insufficient base coverage. Manual inspection of the remaining 9% of the missed calls suggest that either they were also adjacent (but not immediately) to HP regions, or had low coverage resulting in insignificant P-values to pass the IsoPhase SNP call threshold.

**IsoPhase-unique SNP calls are dominantly A to G calls, which suggests RNA editing**

There are 2830 SNP calls unique to IsoPhase, 1776 SNP calls unique to RNA-seq, and 2651 SNP calls unique to genome. We tally the unique calls by Reference to Alternative SNP call in the figure below using the transcribed orientation as the sense strand and show that A to G is the most dominant unique call in IsoPhase and RNA-Seq data. It is not clear why PacBio Iso-Seq is the only platform to call certain SNPs.

Supplementary Note 7 Figure 1: **Percentages of SNP type across those unique in each of PacBio Iso-Seq, RNA-Seq and genome WGS datasets.**

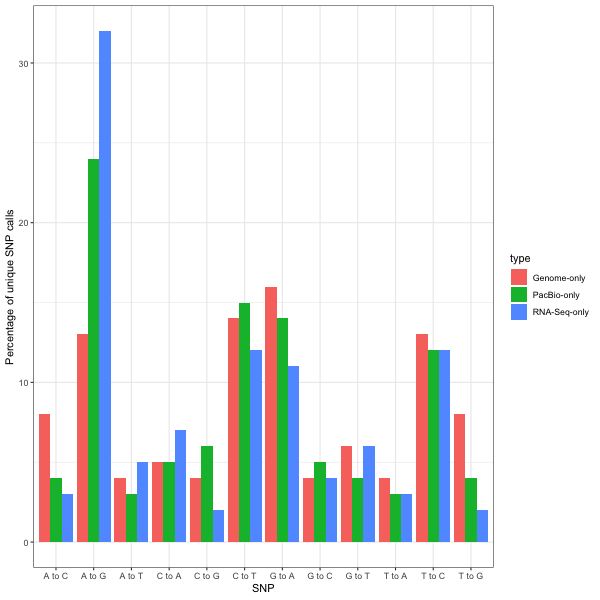

**Rarefaction analysis of covered genes and transcripts**

The subsampling of full-length non-concatamer (FLNC) reads at the level of genes and transcripts showed that brain, kidney and heart datasets were reaching a plateau, suggesting sequencing depth was adequate to discover the majority of transcripts for the annotation process. To ensure saturation of transcripts, additional data was produced for the brain sample, which was run with seven SMRT cells. By extrapolating from the rarefaction curve, 10 SMRT cells per tissue to gather ~1 million FLNC should ensure saturation of transcripts at this developmental stage.

Supplementary Note 7 Figure 2: **Rarefaction analysis of seven tissues in F_1_ animal at the level of** a) gene, b) transcript.

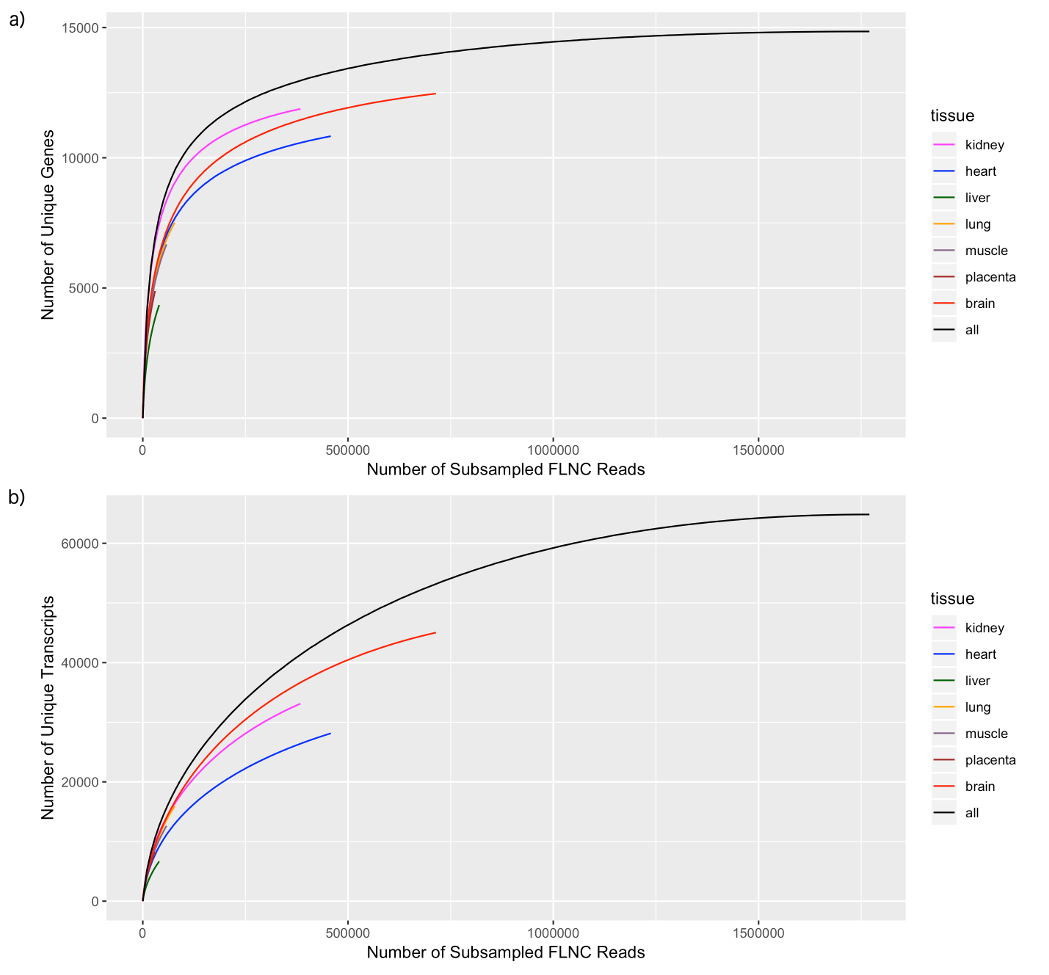
